# De Novo Pyrimidine Synthesis is a Targetable Vulnerability in IDH Mutant Glioma

**DOI:** 10.1101/2021.11.30.470443

**Authors:** Diana D. Shi, Milan R. Savani, Michael M. Levitt, Adam C. Wang, Jennifer E. Endress, Cylaina E. Bird, Joseph Buehler, Sylwia A. Stopka, Michael S. Regan, Yu-Fen Lin, Wenhua Gao, Januka Khanal, Min Xu, Bofu Huang, Rebecca B. Jennings, Dennis M. Bonal, Misty S. Martin-Sandoval, Tammie Dang, Lauren C. Gattie, Sungwoo Lee, John M. Asara, Harley I. Kornblum, Tak W. Mak, Ryan E. Looper, Quang-De Nguyen, Sabina Signoretti, Stefan Gradl, Andreas Sutter, Michael Jeffers, Andreas Janzer, Mark A. Lehrman, Lauren G. Zacharias, Thomas P. Mathews, Timothy E. Richardson, Daniel P. Cahill, Ralph J. DeBerardinis, Keith L. Ligon, Peter Ly, Nathalie Y. R. Agar, Kalil G. Abdullah, Isaac S. Harris, William G. Kaelin, Samuel K. McBrayer

## Abstract

Mutations affecting isocitrate dehydrogenase (IDH) enzymes are prevalent in glioma, leukemia, and other cancers. Although mutant IDH inhibitors are effective against leukemia, they appear less active in aggressive glioma, underscoring the need for alternative treatment strategies. Through a chemical synthetic lethality screen, we discovered that IDH1 mutant glioma cells are hypersensitive to drugs targeting enzymes in the de novo pyrimidine nucleotide synthesis pathway, including dihydroorotate dehydrogenase (DHODH). We developed a genetically engineered mouse model of mutant IDH1-driven astrocytoma and used it and multiple patient-derived models to show that the brain-penetrant DHODH inhibitor BAY 2402234 displays monotherapy efficacy against IDH mutant gliomas. Mechanistically, this vulnerability selectively applies to de novo pyrimidine, but not purine, synthesis because glioma cells engage disparate programs to produce these nucleotide species and because *IDH* oncogenes increase DNA damage upon nucleotide pool imbalance. Our work outlines a tumor-selective, biomarker-guided therapeutic strategy that is poised for clinical translation.

## INTRODUCTION

Glioma is the most common primary malignant brain tumor in adults and one of the most lethal forms of human cancer. Despite intense and continuous efforts to develop novel treatments, no new medical therapies have been approved for adult glioma patients in the last decade. Therefore, there is an urgent unmet clinical need to devise therapeutic strategies that safely and effectively treat these tumors.

Although treatment advances in glioma have been limited, our understanding of the molecular pathogenesis of this disease has grown considerably over the last two decades, driven in large part by the advent of high-throughput sequencing. The discovery of highly recurrent hotspot mutations in the genes encoding isocitrate dehydrogenase (IDH) enzymes, *IDH1* and *IDH2*, provided new insight into the genetic basis of brain tumor formation (Parsons et al., 2008). The presence of an *IDH* mutation is part of the diagnostic criteria for oligodendrogliomas (grades 2–3) and now also defines astrocytoma, IDH-mutant, grades 2–4 (Louis et al., 2021). The vast majority of these tumors are heterozygous for the canonical glioma-associated *IDH1-R132H* mutation (Losman and Kaelin, 2013). IDH1 and IDH2 mutant enzymes gain the neomorphic ability to produce the ‘oncometabolite’ (*R*)-2-hydroxyglutarate [(*R*)-2HG] through a 2-oxoglutarate (2OG)-and NADPH-dependent reaction (Dang et al., 2009). (*R*)-2HG accumulates to millimolar levels in IDH mutant cells and tumors and, due to its structural similarity to 2OG, competitively modulates the activity of 2OG-dependent enzymes (Xu et al., 2011). Notably, inhibition of 2OG-dependent dioxygenases involved in DNA and histone demethylation by (*R*)-2HG profoundly alters chromatin structure, ultimately causing aberrant transcriptional upregulation of the glioma oncogene platelet-derived growth receptor alpha (*PDGFRA*) (Flavahan et al., 2016). (*R*)-2HG also enhances the activity of the EglN1 prolyl hydroxylase, a dioxygenase family member that promotes degradation of the hypoxia-inducible factor 1 alpha (HIF1α) (Koivunen et al., 2012; Tarhonskaya et al., 2014). In preclinical models, HIF1 can suppress glioma cell growth ex vivo and in vivo (Blouw et al., 2003). Together, the cumulative biochemical effects of (*R*)-2HG transform neural cells and initiate glioma formation.

These advances have prompted the design and testing of mutant IDH inhibitors in an effort to block (*R*)-2HG synthesis and to reverse the tumor-promoting effects of this oncometabolite. In contrast to the success of these inhibitors in the treatment of IDH mutant leukemias (Stein et al., 2017), IDH inhibitors have displayed comparably limited antitumor activity against aggressive IDH mutant gliomas in preclinical (Tateishi et al., 2015) and early clinical studies (Mellinghoff et al., 2021). Among patients with recurrent or progressive IDH mutant gliomas, the objective response rates following treatment with an IDH inhibitor in a recent phase I clinical trial were 18% and 0% in patients with non-contrast-enhancing tumors and patients with contrast-enhancing tumors, respectively (Mellinghoff et al., 2021), the latter of which are considered clinically aggressive. These results coincide with cell culture studies demonstrating that a substantial portion of the epigenetic effects of mutant IDH in glial cells are not rapidly reversible and that dependence on (*R*)-2HG synthesis is transient during neural cell transformation (Johannessen et al., 2016; Turcan et al., 2018). Furthermore, there is emerging clinical evidence that a small fraction of IDH mutant gliomas accrue spontaneous copy number alterations affecting *IDH1/2* loci that impair (*R*)-2HG production (Mazor et al., 2017), implying that maintenance of (*R*)-2HG accumulation is not strictly required for IDH mutant glioma growth. Copy number alterations that repress (*R*)-2HG production also occur spontaneously in cultured IDH1 mutant glioma cells and do not reduce cellular fitness (Luchman et al., 2013). Taken together, these data suggest that many IDH mutant gliomas transition toward an (*R*)-2HG-independent phenotype over time, providing rationale to develop alternatives to mutant IDH inhibitors to effectively treat these tumors.

One alternative therapeutic strategy to directly inhibiting IDH mutant oncoproteins would be to exploit the collateral vulnerabilities that these enzymes engender. This approach is conceptually attractive because it leverages the inherent specificity of IDH mutations while uncoupling antitumor efficacy from ongoing (*R*)-2HG dependence. Past work from our group and others supports the feasibility and promise of this strategy. Specific examples include exploiting altered NAD+ metabolism in IDH mutant cells via treatment with NAMPT inhibitors (Tateishi et al., 2015), using PARP inhibitors to target DNA repair liabilities induced by (*R*)-2HG accumulation (Lu et al., 2017; Sulkowski et al., 2017), and blocking compensatory glutamine catabolism caused by (*R*)-2HG-dependent repression of branched chain amino acid catabolism (McBrayer et al., 2018). Identification of these mutant IDH-induced liabilities has directly translated to new treatment strategies for IDH mutant gliomas that are currently being tested in clinical trials.

Because much of the prior research in this area was borne out of hypothesis-driven approaches, we posited that pursuing a complementary, unbiased approach to identifying collateral vulnerabilities induced by IDH mutations might reveal appealing new targets for brain tumor therapy. Considering that metabolic reprogramming is the most proximal consequence of mutant IDH activity in brain tumors and that its biological relevance has been extensively demonstrated and independently validated, we sought to uncover liabilities in cellular metabolism conferred by the *IDH1-R132H* oncogene that could nominate new therapeutic targets in IDH mutant glioma.

## RESULTS

### Mutant IDH1 Sensitizes Cells to De Novo Pyrimidine Synthesis Inhibition

We recently created isogenic IDH1 mutant and wild-type (WT) glioma cell culture models that recapitulate (*R*)-2HG levels in corresponding primary brain tumors (McBrayer et al., 2018). Specifically, we used an endogenous IDH1/2 WT human glioma line, HOG, and infected these cells with a lentivirus encoding the *IDH1-R132H* oncogene or with the empty vector (EV). We used these isogenic models in conjunction with a novel compound screening platform that was developed by two of us (J.E.E. and I.S.H.), named Multifunctional Approach to Pharmacologic Screening (MAPS) (Harris et al., 2019), to comprehensively profile differences in metabolic inhibitor sensitivity between IDH1 mutant and IDH1 WT glioma cells (Figure 1A). This platform uses compound libraries arrayed in 384-well plates to support multiplexed drug sensitivity assays, a plate cytometry instrument that enables high-throughput, imaging-based analysis of cell numbers, and an automated data processing pipeline to calculate integrals of drug dose-response curves. The MAPS platform offers a number of advantages for our experimental objectives relative to conventional pharmacological screening systems. First, we custom-curated a library of inhibitors that collectively target key biochemical pathways, allowing us to deeply probe metabolic dependencies. Second, inhibitors in the MAPS platform are arrayed over 10-point dose series to provide more granular and quantitative data related to drug sensitivity. Third, we incorporated an imaging-based readout of viable cell numbers rather than a metabolic readout of cell viability (i.e., tetrazolium staining or ATP-dependent luminescence) to avoid potentially confounding effects of metabolic perturbations on the latter. We leveraged these unique features of the MAPS platform to identify metabolic vulnerabilities conferred by IDH1 mutations in glioma cells.

**Figure 1.**
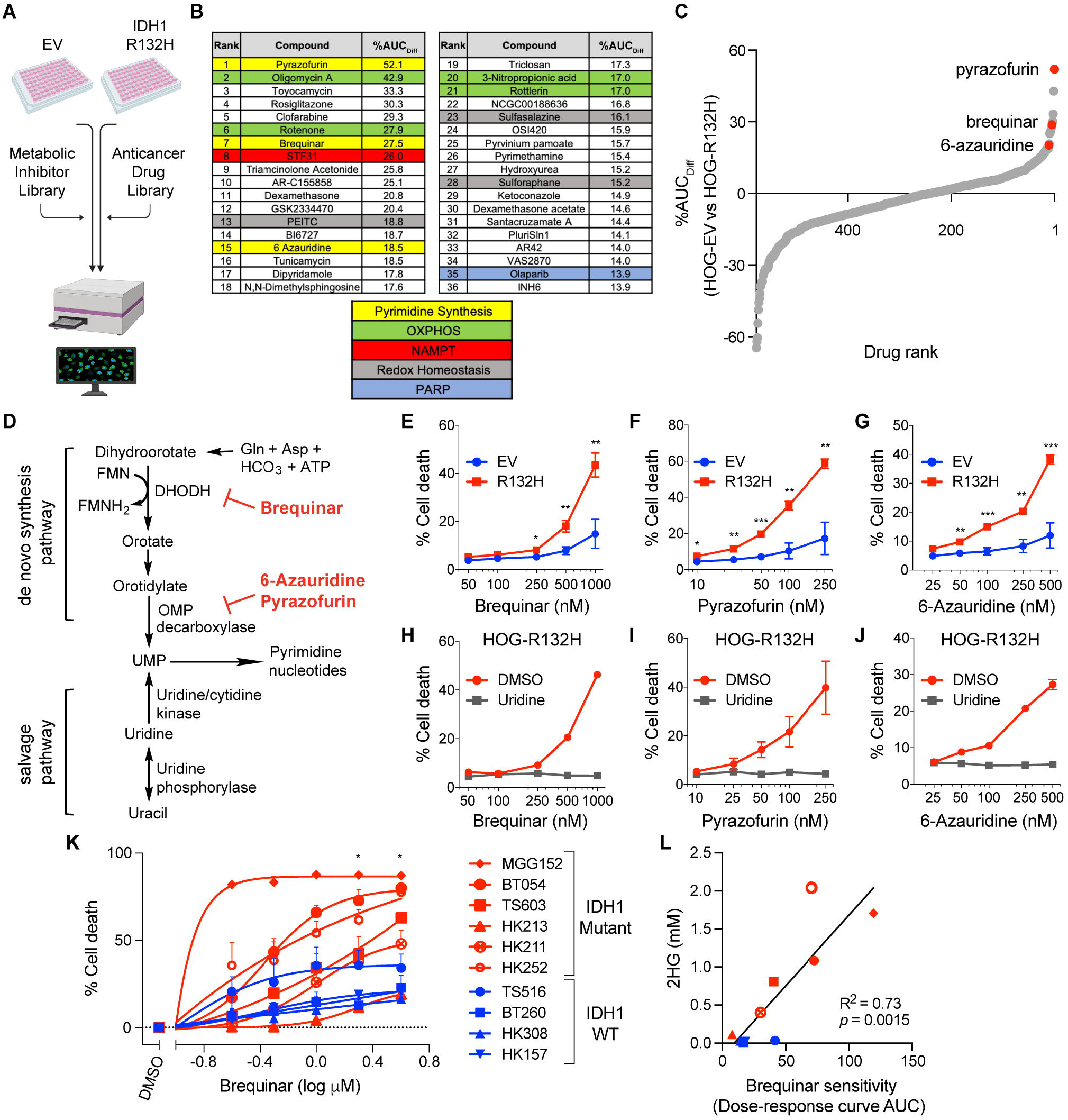
De Novo Pyrimidine Synthesis Inhibitors Preferentially Kill IDH1 Mutant Glioma Cells. (A) Schematic of the Multifunctional Approach to Pharmacologic Screening (MAPS) platform. Isogenic HOG cells expressing either empty vector (EV) or HA-tagged mutant IDH1 (R132H) were treated with 10 different doses of each drug (1 nM to 20 μM) in a metabolic inhibitor library or an anticancer drug library. Cells were stained and counted, with the relative cell number determined at each concentration for both HOG-EV and HOG-R132H cells and used to create area under curve (AUC) values. Percent difference in AUC (%AUC_Diff_) values in HOG-EV and HOG-R132H cells were calculated for all drugs. (B) Top hits from the screen depicted in (A) with the pathways involved shown for subsets of these hits (highlighted and color-coded) (complete list in Supplemental Table 1). Hits were defined as drugs that displayed >10%AUC_Diff_ values, with positive values indicating preferential activity against HOG-R132H cells. In total, 56 hits were identified, and the 36 highest ranking hits are displayed. (C) Waterfall plot of all drugs from both libraries, including three de novo pyrimidine synthesis inhibitors (brequinar, 6-azaurine, and pyrazofurin), based on %AUC_Diff_ values from (A). (D) Schematic showing de novo and salvage pyrimidine synthesis pathways and the enzyme targets of brequinar, 6-azauridine, and pyrazofurin. (E–G) Cell death assays of HOG-EV or HOG-R132H cells treated with brequinar (E), pyrazofurin (F), or 6-azauridine (G) for 48 hours (*n =* 5). (H–J) Rescue experiments using HOG-R132H cells treated with the drugs shown in (E–G) for 48 hours and either 100 μM uridine or DMSO added 24 hours prior (*n* = 3). (K) Cell death assays of 10 patient-derived glioma stem-like (GSC) cell lines (6 IDH1 mutant, 4 IDH1 wild type) treated with BAY 2402234 for two population doublings (PDs) (*n* ≥ 3). Data are normalized to set cell death = 0% in DMSO-treated cells. (L) Correlation between 2HG levels [measured by gas chromatography-mass spectrometry (GC-MS)] and sensitivity of GSC lines to brequinar (red: IDH mutant; blue: IDH1 WT) as in (K). Symbols representing individual cell lines are the same as in (K). For all panels, data presented are means ± SEM; \**p* < .05, \*\**p* < .01, \*\*\**p* < .001. For (L), *p*-value was determined by simple linear regression analysis. For all other parts, two-tailed *p*-values were determined by unpaired *t*-test.

IDH1 mutant (HOG-R132H) and IDH1 WT (HOG-EV) isogenic HOG stable cell lines were screened against our custom metabolic inhibitor library as well as a commercially available anticancer drug library. In sum, we screened 546 unique compounds that target a wide range of cellular pathways and processes (Figure S1A). We identified multiple compounds that selectively reduced the fitness of IDH1 mutant cells versus IDH1 WT cells (Figures S1B and S1C). Many of these compounds, hereafter referred to as ‘hits’, target enzymes involved in mitochondrial, lipid, or nucleotide metabolism (Figure S1D). Taken together, these data indicate that the *IDH1-R132H* oncogene induces liabilities related to cellular metabolism.

Among the hits were compounds targeting pathways and enzymes previously described to represent dependencies conferred by IDH mutations (Figure 1B and Table S1). These compounds include NAMPT, PARP, and oxidative phosphorylation inhibitors, as well as agents that promote oxidative stress (Grassian et al., 2014; Lu et al., 2017; McBrayer et al., 2018; Sulkowski et al., 2017; Tateishi et al., 2015). These findings underscore the utility of the MAPS platform in identifying oncogene-driven dependencies. Interestingly, one unanticipated class of drugs that scored strongly in our screen was de novo pyrimidine synthesis inhibitors. Indeed, three of the top twenty compounds that displayed the greatest selectivity for IDH1 mutant glioma cells belong to this drug class (Figures 1B–D, and S1E–G). This effect appeared to be specific because inhibitors of purine metabolism, including the de novo purine synthesis inhibitor lometrexol, did not reduce cell fitness in a mutant IDH-dependent manner (Figures S1H and S1I).

To attempt to validate our findings from the screen, we assessed the apoptotic response of HOG-EV and HOG-R132H stable lines to treatment with three inhibitors of de novo pyrimidine synthesis: the dihydroorotate dehydrogenase (DHODH) inhibitor brequinar and the orotidylate monophosphate (OMP) decarboxylase inhibitors pyrazofurin and 6-azauridine. In agreement with our screening data, all three of these inhibitors preferentially killed IDH1 mutant glioma cells (Figures 1E–G). The cytotoxic effects of pyrimidine synthesis inhibitors in HOG-R132H cells were on-target because they could be fully rescued by stimulating pyrimidine nucleotide salvage with supraphysiological levels of uridine (Figures 1H–J).

We next asked whether the DHODH inhibitor brequinar preferentially kills patient-derived glioma stem-like cell (GSC) lines carrying endogenous *IDH1-R132H* mutations. We collected a panel of ten GSC lines, six of which were heterozygous for the *IDH1-R132H* mutation and four of which were *IDH1* WT. Each of these lines was treated with a range of doses of brequinar for two population doublings (Figure S1J) and cell viability was subsequently analyzed. Brequinar treatment induced greater rates of cell killing in the IDH1 mutant GSC lines relative to their IDH1 WT counterparts, with the exception of one line, HK213 (Figure 1K). Further examination revealed that within our panel of GSC lines, 2HG content correlated closely with brequinar sensitivity (Figure 1L). Notably, HK213 cells displayed the lowest 2HG content within the IDH1 mutant GSCs, possibly explaining their insensitivity to this drug (Figure 1K and L). Collectively, our studies using an engineered, isogenic cell culture model and patient-derived GSC lines implicate de novo pyrimidine synthesis as an unappreciated vulnerability induced by the *IDH1-R132H* oncogene in glioma.

### IDH1 Mutant Gliomas are Sensitive to the Brain-Penetrant DHODH Inhibitor BAY 2402234

To translate these findings, we next sought to identify a brain-penetrant inhibitor of de novo pyrimidine synthesis that could be used to target this pathway in IDH1 mutant gliomas in vivo. Many classical inhibitors of this pathway are nucleoside analogues with poor CNS penetration. Therefore, we investigated a newly developed DHODH inhibitor, BAY 2402234, that was tested in a clinical trial for treating leukemia (NCT03404726) (Christian et al., 2019). To identify a metabolic biomarker of DHODH inhibition in peripheral tissues that could be applied to pharmacodynamic studies of BAY 2402234 in brain tissues, we treated non-tumor-bearing mice with BAY 2402234 or vehicle and performed unbiased metabolite profiling on heart and liver samples. We found that the product of DHODH (orotate) and an intermediate in the de novo pyrimidine synthesis pathway upstream of DHODH (carbamoyl aspartate) were decreased and increased, respectively, in response to BAY 2402234 (Figures 2A and B). Therefore, we used the ratio of orotate to carbamoyl aspartate as a biomarker of DHODH inhibition in vivo. Importantly, the orotate to carbamoyl aspartate ratio was also suppressed in the brain tissue of non-tumor-bearing mice following BAY 2402234 treatment (Figure 2C), indicating that this agent is brain penetrant and effectively engages DHODH in the CNS. Moreover, daily BAY 2402234 administration was well tolerated over the course of a 2-week treatment (Figure 2D). Complementary in vitro studies revealed that BAY 2402234 preferentially killed IDH1 mutant HOG cells and GSC lines in a manner that was uridine-dependent and correlated with intracellular 2HG content, thereby recapitulating the effects of the structurally distinct DHODH inhibitor brequinar (Figures 2E–2H). Based on these findings, we proceeded to test the antitumor activity of BAY 2402234 in mouse models of IDH1 mutant glioma.

**Figure 2.**
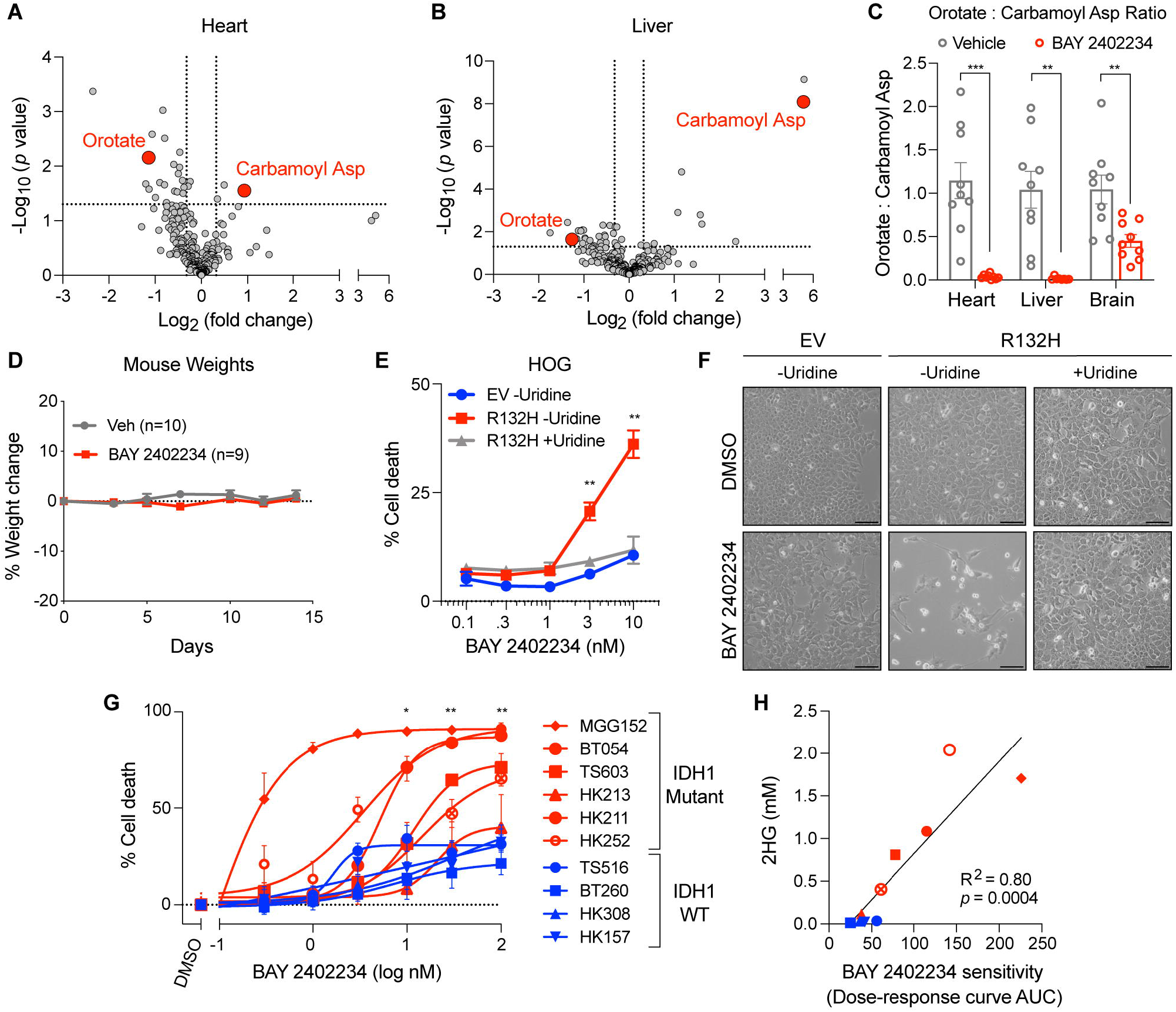
The DHODH Inhibitor BAY 2402234 is Brain-Penetrant in Mice and Selectively Kills IDH1 Mutant Glioma Cells. (A and B) Volcano plot of metabolites [measured by liquid chromatography-tandem mass spectrometry (LC-MS/MS)] in the heart (A) and liver (B) of ICR SCID mice treated with BAY 2402234 (4 mg/kg PO QD) relative to vehicle for 3 days (*n =* 9 per cohort). Carbamoyl asp = carbamoyl aspartate. (C) Ratio of orotate to carbamoyl aspartate in the heart, liver, and brain of mice treated with BAY 2402234 or vehicle as in (A) (*n =* 9 per cohort). (D) Change in weight over time of mice treated with either BAY 2402234 (4 mg/kg PO QD) or vehicle for 14 days. (E) Cell death assays of HOG-EV or HOG-R132H cells treated with BAY 2402234 at the indicated concentrations for 72 hours (*n* = 3). Where indicated, 100 μM uridine was added 24 hours before BAY 2402234. (F) Representative photomicrographs of cells in (E). Scale bars = 100 μm. (G) Cell death assays of human GSC lines treated with BAY 2402234 or DMSO for two PDs (*n* ≥ 3). Data are normalized to set cell death = 0% in DMSO-treated cells. (H) Correlation between 2HG levels (measured by GC-MS) and sensitivity of GSC lines to BAY 2402234 (red: IDH mutant; blue: IDH WT) as in (G). Symbols representing individual cell lines are the same as in (G). For all panels, data presented are means ± SEM; \**p* < .05, \*\**p* < .01, \*\*\**p* < .001. For (H), *p*-value was determined by simple linear regression analysis. For all other parts, two-tailed *p*-values were determined by unpaired *t*-test.

For initial in vivo efficacy studies of BAY 2402234, we used the MGG152 orthotopic xenograft glioma model, which was derived from a recurrent IDH1 mutant grade 4 glioma and is resistant to mutant IDH1 inhibitor treatment (Tateishi et al., 2015; Wakimoto et al., 2014). Treating MGG152 tumor-bearing mice with BAY 2402234 depleted orotate and increased carbamoyl aspartate pervasively throughout tumor tissue, as visualized by matrix-assisted laser desorption ionization mass spectrometry imaging (MALDI-MSI) (Figure 3A), demonstrating effective target engagement. Furthermore, BAY 2402234 accumulated to ∼150 nM in tumor tissue following oral administration (Figure 3B), which exceeded the concentration of this drug required to kill IDH1 mutant glioma cells in culture (Figures 2E and 2G). BAY 2402234 treatment prolonged survival relative to vehicle-treated control mice (Figure 3C) and the magnitude of the survival benefit was comparable to, albeit slightly less than, the effect of radiotherapy (Figure 3D), which is a cornerstone of the standard-of-care treatment regimen for glioma patients. This effect was specific because BAY 2402234 did not prolong the survival of mice bearing orthotopic tumors established from the TS516 IDH1/2 WT GSC line (Figure 3E). Importantly, therapy failure in this IDH WT glioma model cannot be explained by poor target engagement, as the orotate to carbamoyl aspartate ratio decreased similarly upon BAY 2402234 treatment in MGG152 and TS516 tumors (Figures 3F and G). These data support the ability of BAY 2402234 to effectively penetrate brain tumor tissue and show that MGG152 grade 4 IDH1 mutant gliomas display an ongoing requirement for DHODH activity.

**Figure 3.**
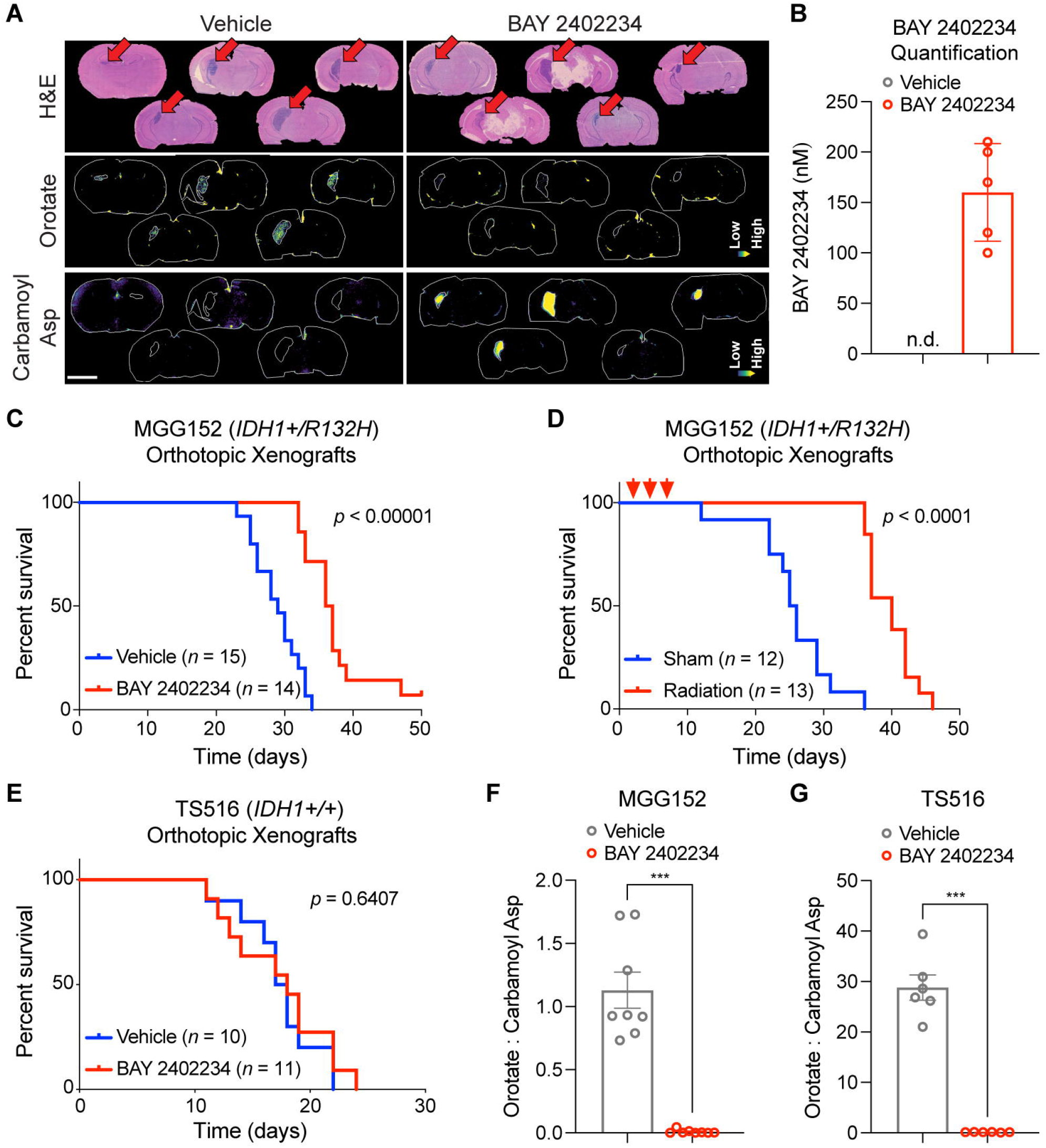
Treatment with BAY 2402234 Improves Survival of Mice Bearing IDH1 Mutant Glioma Orthotopic Xenografts. (A) Representative brain tissue sections from mice bearing MGG152 (IDH1 mutant) orthotopic glioma xenografts and treated with BAY 2402234 (4 mg/kg PO QD) or vehicle for 3 days (*n* = 5 per cohort). Brains were harvested 4 hours after the final dose. Top panels: H&E staining showing site of tumor development. Middle and bottom panels: relative orotate (middle) and carbamoyl aspartate (bottom) levels throughout the whole brain as determined by MALDI-MSI. Scale bar = 3 mm. (B) MALDI-MSI-based absolute quantification of BAY 2402234 accumulation in tumor tissues in (A). n.d. = not detected. (C and D) Kaplan-Meier survival curves of mice bearing MGG152 orthotopic xenografts treated with BAY 2402234 (4 mg/kg PO QD) or vehicle (C) or cranial radiation (9 Gy in 3 fractions) or sham (D). BAY 2402234 and vehicle treatments were administered continuously starting Day 12 after tumor cell injection (displayed as Day 0 in C and D). Red arrows in (D) indicate timing of radiation doses. (E) Kaplan-Meier survival curves of mice bearing TS516 (IDH WT) orthotopic glioma xenografts treated with BAY 2402234 or vehicle. Treatments were the same as in (C), except they began Day 9 after tumor cell injection (displayed as Day 0 in E). (F and G) Orotate to carbamoyl aspartate ratios in tumor tissues from MGG152 (*n =* 8 per cohort) (F) and TS516 (*n =* 6 per cohort) (G) xenografts that were treated as in (A). Metabolites were measured by LC-MS/MS (F) or liquid chromatography-mass spectrometry (LC-MS) (G). For all panels, data presented are means ± SEM; \**p* < .05, \*\**p* < .01, \*\*\**p* < .001. In (C–E), *p*-values were calculated by log-rank test and all studies were repeated twice. In (F) and (G), two-tailed *p*-values were determined by unpaired *t*-test.

### Creation of a Genetically Engineered Mouse Model of Mutant IDH1-Driven Astrocytoma

One general concern regarding patient-derived xenograft models of glioma is that they have largely been created from highly proliferative gliomas, whereas most IDH mutated brain tumors are relatively indolent, with smaller proportions of mitotic cells. This concern raised the question of whether more indolent brain tumors with *IDH* mutations would likewise respond to DHODH inhibitors. This question is important because nucleotide synthesis dependencies in other cancers have principally been associated with rapidly proliferating cells. To address this issue, we sought to complement our efficacy studies using the highly proliferative MGG152 glioma model by developing a genetically engineered mouse (GEM) model of mutant IDH1-driven grade 3 astrocytoma and using it to test BAY 2402234 therapy.

Brain-specific activation of the *IDH1-R132H* oncogene by itself is not sufficient to promote gliomagenesis in mice (Bardella et al., 2016; Sasaki et al., 2012a). We hypothesized that more fully recreating the mutational landscape of astrocytoma, IDH-mutant, grade 3 in the brain tissue of adult mice would suffice to cause formation of tumors that faithfully recapitulate this disease and isolate the contributions of specific co-mutation drivers. To identify mutations that frequently co-occur with *IDH1* mutations in lower grade gliomas, we analyzed the TCGA lower grade glioma clinical genomics dataset and identified a subset of astrocytomas that harbored concurrent mutations in *IDH1*, *TP53*, and *ATRX* genes along with alterations affecting *PIK3CA* or *PIK3R1* genes, which encode the two subunits of PI3-kinase (PI3K) (Figure 4A). Some *PIK3CA* and *PIK3R1* mutations co-occurred with both *IDH1* mutations and *CIC* mutations, with the latter being a hallmark of oligodendrogliomas. Statistical analysis of co-occurrence between *IDH1* mutations and those in *TP53* or *ATRX* revealed highly significant positive associations in the setting of both lower grade gliomas and in grade 4 gliomas (Table S2). *PIK3R1* mutations were associated with *IDH1* mutations in grade 4 gliomas and trended toward significant co-occurrence in lower-grade gliomas, suggesting that this might reflect an important pathway of IDH1 mutant glioma progression. Mutations in *PIK3CA* were observed in both *IDH1* mutated and WT tumors independent of grade. Although PTEN is a negative regulator of PIK3CA, *PTEN* mutations appeared to be mutually exclusive with *IDH1* mutations. We next asked whether PI3K mutations define a subset of IDH1 mutant lower grade glioma patients with particularly poor outcomes, a finding that would be consistent with cooperation between these oncogenes. Indeed, patients with coincident *IDH1* and *PIK3CA* and/or *PIK3R1* mutations displayed inferior survival compared with those that harbored an *IDH1* mutation but lacked *PIK3CA* and *PIK3R1* mutations (Figure 4B). In support of these findings, a recent study identified mutation of the *PIK3R1* gene as an independent negative prognostic indicator in a cohort of *IDH1* mutated astrocytoma patients (Aoki et al., 2018).

**Figure 4.**
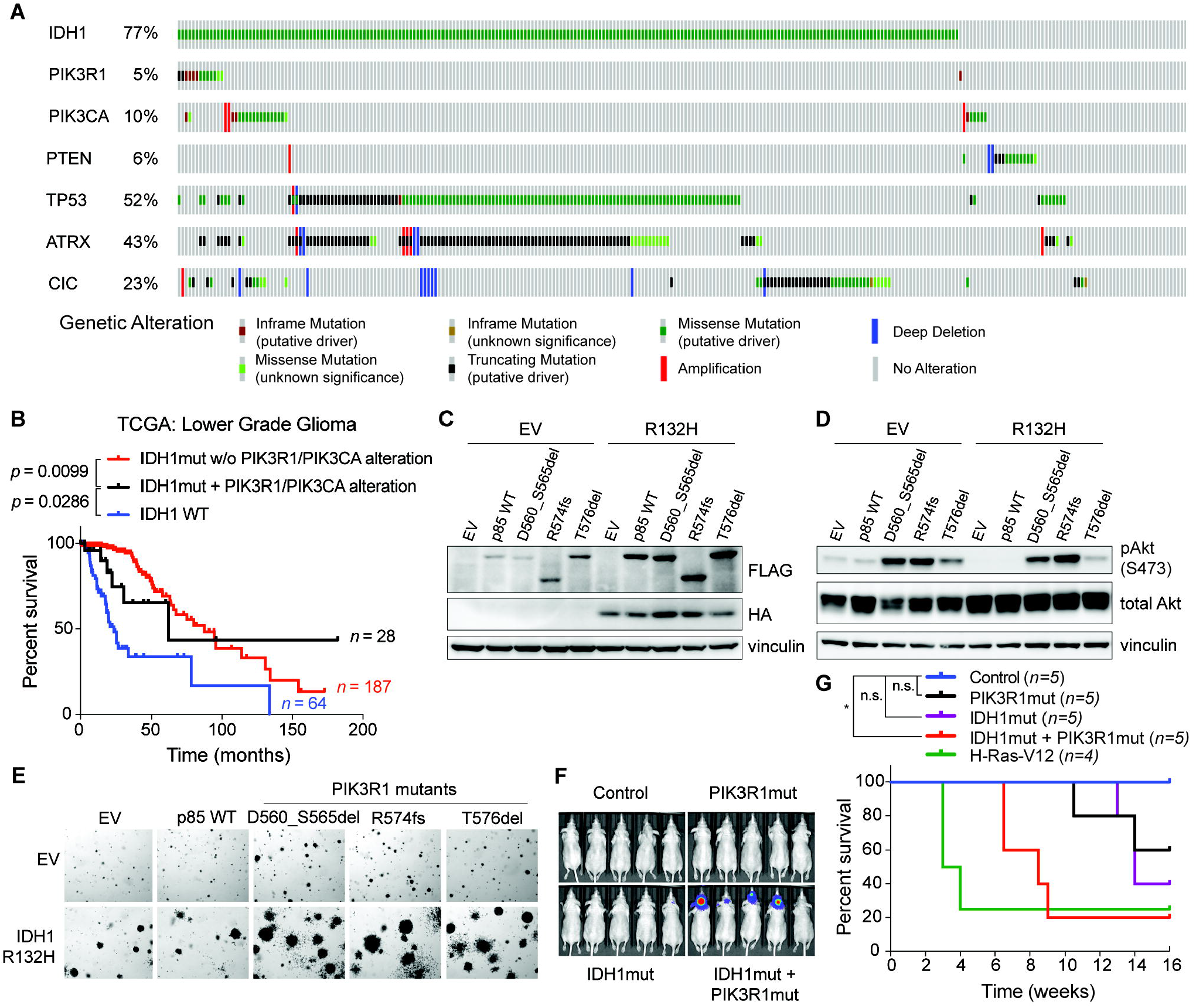
*IDH1* and *PIK3R1* Oncogenes Cooperate to Transform Immortalized Astrocytes. (A) Oncoprint of samples in the Brain Lower Grade Glioma TCGA dataset showing frequency and co-occurrence of mutations in *IDH1*, *PIK3R1, PIK3CA, PTEN, TP53*, *ATRX*, and *CIC*. (B) Kaplan-Meier survival curves of human glioma patients with the indicated genotypes. (C and D) Immunoblot analysis of human astrocytes (immortalized with hTERT, E6, and E7) cultured under low (0.5%) fetal bovine serum conditions and infected to express HA-tagged IDH1-R132H [or with the empty vector (EV)] and then superinfected to express FLAG-tagged wild-type or mutant (D560_S565del, R574fs, or T576del) p85 (protein product of *PIK3R1* gene) or the corresponding EV. All cells were also infected to express firefly luciferase and GFP. PI3K phosphorylates Akt S473. (E) Soft agar colonies formed by the astrocyte cell lines used in (C) and (D) under standard serum conditions. *n* = 5, representative cultures are shown. (F and G) Bioluminescence imaging (F) and Kaplan-Meier curves (G) of nude mice that were orthotopically injected with astrocytes expressing EV/EV (control), IDH1^R132H^/EV (IDH1mut), EV/p85^D560_S565del^ (PIK3R1mut), IDH1^R132H^/p85^D560_S565del^ (IDH1mut + PIK3R1mut), or H-Ras-V12 (as a positive control). Bioluminescence imaging in (F) was performed 6 weeks post-cell implantations. Only 1 mouse implanted with H-Ras-V12-expressing NHA cells survived past Week 4; this cohort was therefore not imaged at Week 6. \**p* < .05, n.s. = not significant. *p*-values were determined by log-rank test.

We hypothesized that mutating *Idh1* with *Pik3r1* or *Pik3ca* in neural cells in the setting of astrocytoma-specific *Atrx* and *Trp53* mutations would promote the development of gliomas in mice within a timeframe suitable for preclinical studies. As a risk-assessment study, we first determined whether *PIK3R1* and *IDH1* oncogenes cooperate to transform astrocytes (NHA cells) that were immortalized via expression of HPV E6 (which phenocopies *TP53* mutations) and E7 (which phenocopies *RB* mutations) proteins as well as hTERT (which phenocopies ATRX loss). Using NHA cells that were previously infected to express the *IDH1-R132H* oncogene or with an empty vector and subsequently passaged >25 times (Koivunen et al., 2012), we co-expressed glioma patient-derived *PIK3R1* mutants (Quayle et al., 2012), a *PIK3R1* WT cDNA, or the empty vector (Figure 4C). After validating that the *PIK3R1* mutants activate PI3K signaling (Figure 4D), we tested this panel of cell lines in in vitro and in vivo transformation assays. In agreement with our hypothesis, we found that *PIK3R1* and *IDH1* mutations cooperate to promote anchorage-independent growth (Figures 4E and S2A) and orthotopic tumor formation in immunodeficient mice (Figures 4F and G; Figures S2B and C).

We sought to apply these findings to create a GEM model of mutant IDH1-driven grade 3 astrocytoma by combining recombinant adeno-associated virus (AAV), CRISPR/Cas9, and transgenic mouse technologies. First, we bred four compound transgenic mice featuring the following alleles in various combinations: *LSL-Idh1-R132H*, *LSL-Pik3ca-H1047R*, and *LSL-Cas9* (Figure 5A) (Adams et al., 2011; Platt et al., 2014; Sasaki et al., 2012a). Next, we made AAVs expressing a Cre cDNA and sgRNAs targeting *Trp53* and *Atrx* genes. Finally, we injected these AAVs into the subventricular zone of the brains of adult compound transgenic mice to induce loss-of-function mutations in *Trp53* and *Atrx* in the presence or absence of gain-of-function *Idh1* and/or *Pik3ca* mutations. Hereafter, these compound transgenic mice will be referred to by abbreviations summarizing their distinct transgenic profiles: *LSL-**P**ik3ca^H1047R/+^*;***I****dh1^LSL-R132H/+^*;*LSL-**C**as9^+/-^* (***PIC*** cohort), *2) **I**dh1^LSL-R132H/+^*;*LSL-**C**as9^+/-^* (***IC*** cohort), 3) *LSL-**P**ik3ca^H1047R/+^*;*LSL-**C**as9^+/-^* (***PC*** cohort), and 4) *LSL-**C**as9^+/-^* (***C*** cohort). In accordance with predictions from our studies involving engineered NHA cells in Figure 4, we observed glioma formation by magnetic resonance imaging (MRI) in AAV-injected ***PIC*** mice starting 10 months after virus injection (Figure 5B). We validated that these gliomas displayed 2HG upregulation, recombination of *Pik3ca* and *Idh1* mutant alleles, and inactivating mutations in *Trp53* and *Atrx* genes (Figures S3A–F). Unexpectedly, many AAV-injected ***PC*** and ***PIC*** mice developed sarcomas at the site of virus injection in the skull that contributed to mortality in these cohorts (Figure 5C and Figures S3G). However, sarcoma-free, AAV-injected ***PIC*** mice went on to develop grade 3 astrocytomas that were either not (***IC*** and ***C***) or very infrequently (***PC***) observed in other cohorts (Figure 5D). These murine gliomas histologically resembled their human counterparts, exhibiting a diffuse, invasive morphology, and expression of the glial lineage markers Gfap and Olig2 (Figures 5E and 5F). Taken together, these data show that oncogenic *Idh1*, *Pik3ca*, *Trp53*, and *Atrx* mutations cooperate to cause autochthonous gliomas to form in mice that resemble astrocytoma, IDH-mutant, grade 3 in human patients.

**Figure 5.**
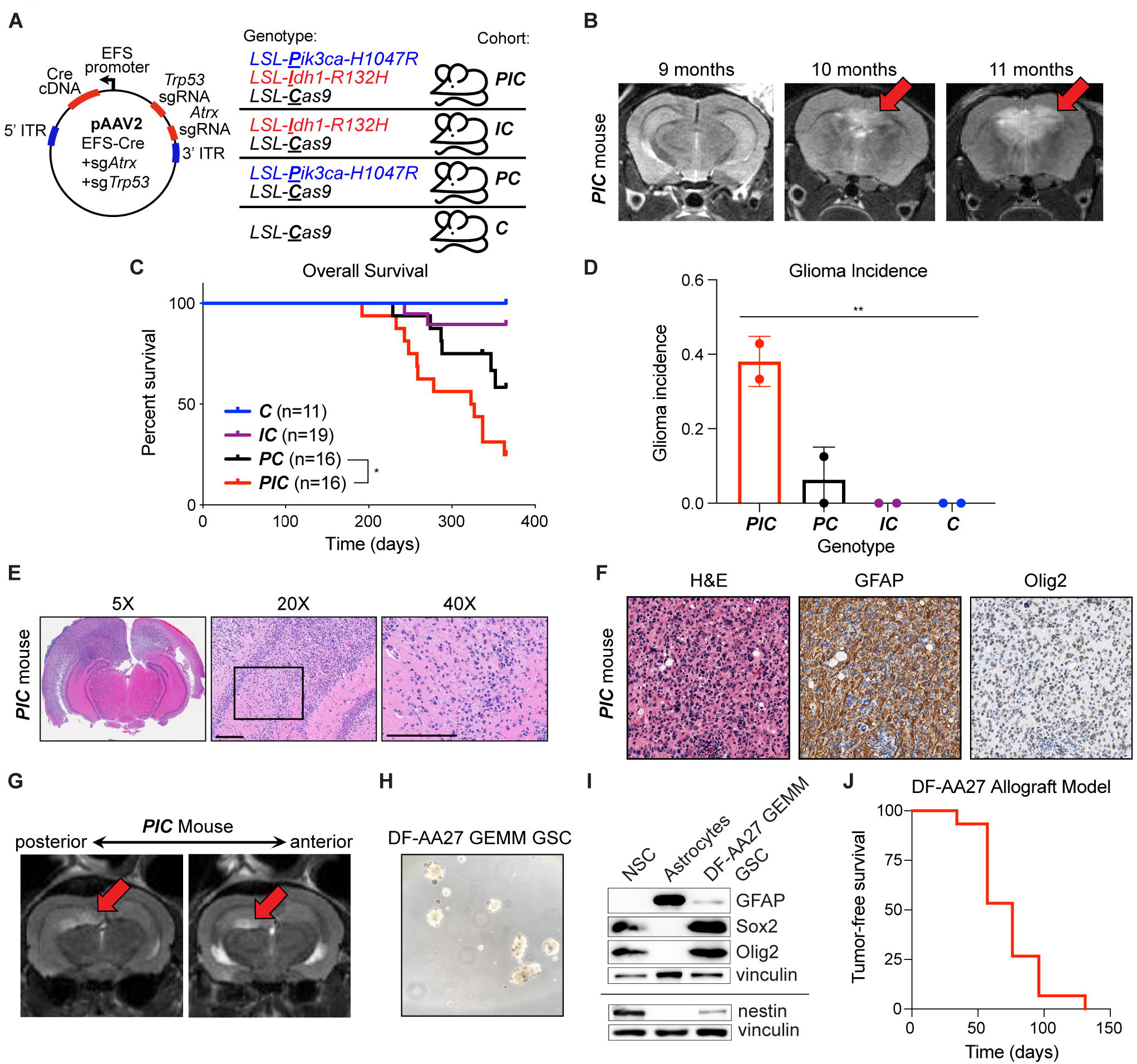
Creation of Genetically Engineered Mouse Models of Lower-Grade Astrocytoma. (A) Mouse injection scheme. An adeno-associated virus (AAV) encoding Cre-recombinase and sgRNAs targeting murine *Atrx* and *Trp53* genes was intracranially injected into four different engineered mouse strains (***PIC***, ***IC***, ***PC***, and ***C***) as shown, where the indicated genes were placed downstream of a *loxP*-stop-*loxP* (LSL) cassette. Mice were monitored for tumor initiation with serial monthly MRI scans. (B) Representative MRI images of one mouse from the ***PIC*** cohort at the indicated time points after intracranial AAV injection, as in (A). (C) Kaplan-Meier survival curves of AAV-injected mice, as in (A) (date of injection = time 0). (D) Cumulative glioma incidence in mice (as determined by MRI, histopathologic analysis, or both), as in (A), in the 12 months following AAV injection. Glioma incidence was measured in two separate cohorts of mice; mice that developed injection-site sarcomas were censored. (E and F) Hematoxylin and eosin (H&E) (E) and immunohistochemical (IHC) (F) stained sections of the brain from a representative AAV-injected ***PIC*** mouse. Higher magnification images showing pathologic areas of infiltrative growth at the interface of tumor and normal brain parenchyma. In (E), black box in middle panel indicates region in right panel; scale bars = 200 μm. (G) Representative MRI images from an AAV-injected ***PIC*** mouse with arrows indicating areas of tumor growth, which was harvested and cultured in 5% oxygen to generate a neurosphere-forming GSC line (DF-AA27 GEMM GSC). (H and I) Photomicrograph (H) and immunoblot analysis (I) of DF-AA27 GEMM GSCs, human neural stem cells (NSC), and primary human astrocytes. (J) Kaplan-Meier tumor-free survival curve of ICR SCID mice that were orthotopically injected with DF-AA27 cells (date of injection = time 0) (*n =*15). Mice were monitored for tumor formation with serial MRI scans. For all panels, data presented are means ± standard deviation; \**p* < .05, \*\**p* < .01. In (C), *p*-value was determined by log-rank test. In (D), *p*-value was determined by one-way ANOVA.

### DHODH Inhibition Displays Monotherapy Activity in Grades 3 And 4 IDH Mutant Gliomas

To circumvent issues related to sarcoma formation and protracted tumor latency, we derived a permanent and transplantable GSC line from an astrocytoma that formed in an AAV-injected ***PIC*** mouse (Figures 5G and 5H). This GSC line, which we termed the DF-AA27 GEMM GSC model, grew as neurospheres, expressed appropriate lineage and stemness (Gfap, Olig2, Sox2, Nestin) markers, displayed elevated 2HG levels that were depleted by the mutant IDH1 inhibitor AGI-5198, and harbored engineered mutations in *Idh1*, *Pik3ca*, *Trp53*, and *Atrx* genes (Figures 5H and I; Figures S3H–L). We next performed secondary transplants of DF-AA27 GEMM GSCs into the brains of recipient immunodeficient mice and found that these cells formed allografts within one to four months (Figure 5J; Figure S3M) that retained high 2HG levels (Figure S3N). Therefore, this DF-AA27 GEMM-derived allograft model represents an appealing platform to evaluate experimental therapies in lower grade gliomas.

After establishing that the DF-AA27 GEMM GSC line exhibits similar sensitivity to BAY 2402234 as patient-derived IDH1 mutant human GSC lines in vitro (Figures 2G and 6A), we proceeded to test the efficacy of this drug in the DF-AA27 GEMM-derived orthotopic allograft model of astrocytoma. BAY 2402234 treatment reduced the orotate to carbamoyl aspartate ratio in allografts and markedly attenuated tumor growth (Figures 6B–D). Of the eight DF-AA27 allografts treated with BAY 2402234, three tumors regressed and another three displayed reduced growth relative to vehicle-treated controls (Figure 6C). These data suggest that de novo pyrimidine synthesis represents a novel oncogene-induced metabolic vulnerability in IDH1 mutant gliomas that manifests in both grade 3 and grade 4 gliomas. To further test this hypothesis, we created a panel of patient-derived glioma organoid models from primary surgical tissue specimens (Abdullah et al., 2021). This panel featured five organoids derived from glioblastoma, IDH-wildtype samples and three IDH1 mutant glioma organoids: one derived from an astrocytoma, IDH-mutant, grade 4 sample, one from an oligodendroglioma, IDH-mutant, grade 3 sample, and one from an astrocytoma, IDH-mutant, grade 3 sample. After confirming that these organoids maintain histological features of the respective parental tumors, we treated them with BAY 2402234 or DMSO and measured apoptosis induction via immunohistochemical quantification of cells expressing cleaved caspase 3 (CC3) (Figure 6E). All three IDH1 mutant organoid models tested showed increased incidence of CC3-expressing cells upon BAY 2402234 treatment, whereas only one of the five IDH1/2 WT glioma organoids responded similarly. These data support our hypothesis that DHODH hyperdependence in IDH1 mutant gliomas is not confined to highly proliferative grade 4 tumors.

**Figure 6.**
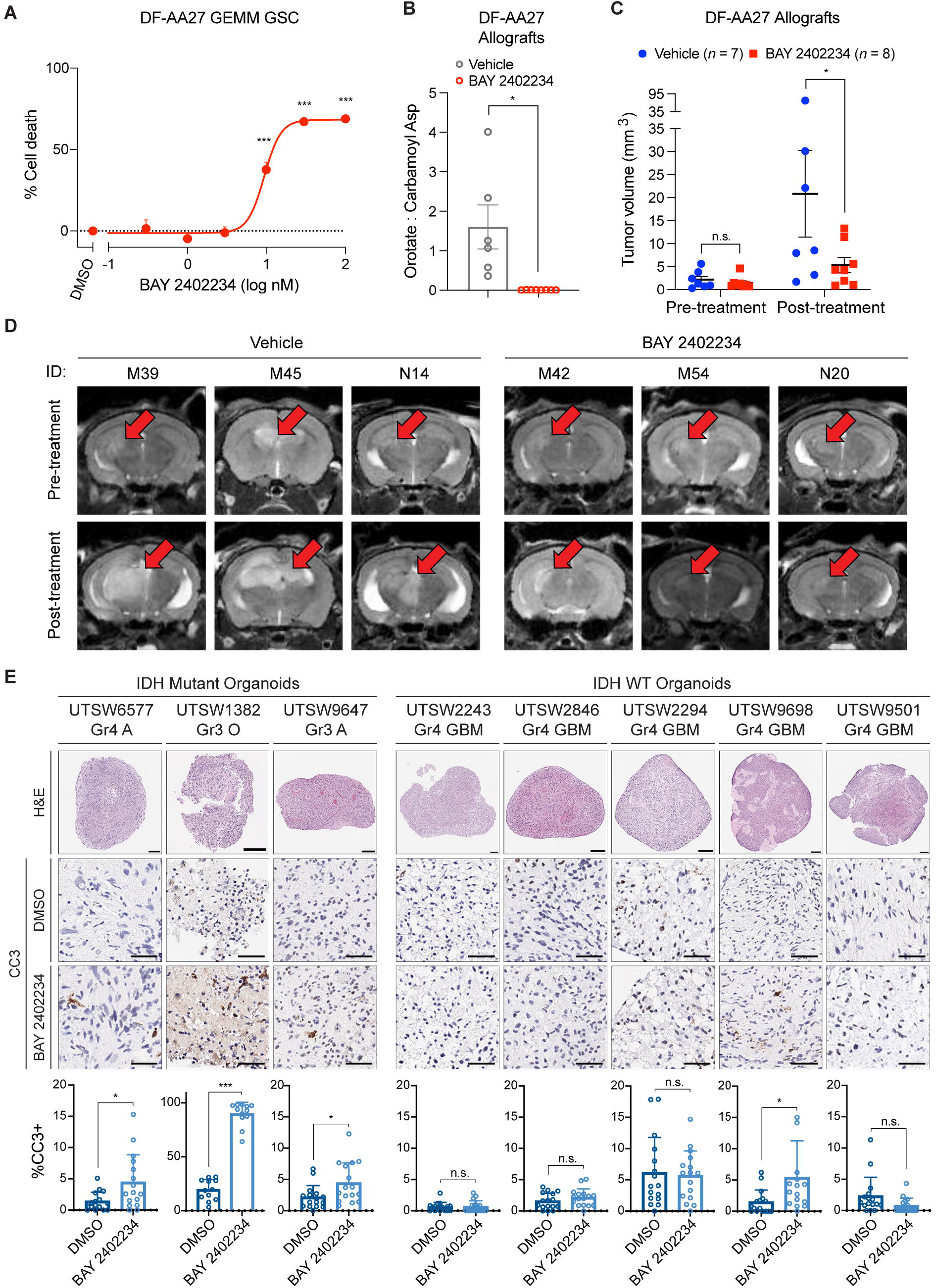
Organoid and Genetically Engineered Allograft Models of IDH Mutant Glioma Respond to DHODH Inhibition. (A) Cell death assay in DF-AA27 GEMM GSCs treated with BAY 2402234 or DMSO for 4 days (*n =* 4). Data are normalized to set cell death = 0% in DMSO-treated cells. (B) Orotate to carbamoyl aspartate ratios (measured by LC-MS) in DF-AA27 orthotopic allografts in mice treated with BAY 2402234 (4 mg/kg PO QD) or vehicle for 3 days (*n =* 6 per cohort). Allograft tissues were harvested 4 hours after the final dose. (C and D) Tumor volume measurements based on MRI images (C) and representative MRI images (D) of mice with DF-AA27 orthotopic allografts treated continuously with BAY 2402234 (4 mg/kg PO QD) or vehicle for 3 weeks following tumor formation, as detected by MRI. Pre-treatment and post-treatment MRIs are at the time of enrollment and 3 weeks later, respectively. (E) Histology and IHC detection and quantification of cleaved caspase 3 (CC3) levels in patient-derived glioma organoids treated with 30 nM BAY 2402234 or DMSO for 14 days. Three IDH mutant and five IDH WT glioma organoid models were evaluated. Top row: H&E staining and gross appearance of DMSO-treated organoids. Scale bars = 200 μm. Middle and bottom rows: IHC staining for CC3 after treatment. Scale bars = 50 μm. Bar graphs: quantification of the percentage of CC3-positive cells in organoids after treatment. Gr = tumor grade, A = astrocytoma, O = oligodendroglioma, GBM = glioblastoma; “UTSW” labels identify organoid models. For (A–C), data presented are means ± SEM. For (E), data presented are means ± standard deviation. \**p* < .05, \*\**p* < .01, \*\*\**p* < .001, n.s. = not significant. Two-tailed *p*-values were determined by unpaired *t*-test.

### Mechanisms of De Novo Pyrimidine Synthesis Hyperdependence in IDH Mutant Glioma

These findings prompted us to investigate two key mechanistic questions. First, why do IDH1 mutant glioma cells rely on de novo pyrimidine, but not de novo purine, synthesis to maintain viability? Second, how do IDH1 mutations cause ongoing dependence on de novo pyrimidine synthesis for glioma cell viability?

To address our first question, we cultured a patient-derived IDH1 mutant oligodendroglioma grade 3 GSC line, BT054, in human plasma-like medium (HPLM) (Cantor et al., 2017) to directly compare the metabolic effects of de novo pyrimidine and de novo purine synthesis inhibition under physiologically relevant conditions. We first confirmed that BAY 2402234 and lometrexol, an inhibitor of glycinamide ribonucleotide transformylase (GART) in the de novo purine synthesis pathway, engaged their respective enzyme targets. BAY 2402234 exposure increased levels of dihydroorotate, the substrate of DHODH, while lometrexol treatment increased levels of glycinamide ribonucleotide (GAR), the substrate of GART (Figure 7A). Next, we quantified representative pyrimidine (UMP) and purine nucleotides (AMP) and found that BAY 2402234 potently reduced pyrimidine nucleotide synthesis, whereas lometrexol did not affect purine nucleotide synthesis. These data suggest that IDH1 mutant glioma cells rely predominantly on the de novo pathway for pyrimidine synthesis and the salvage pathway for purine synthesis (Figure 7B).

**Figure 7.**
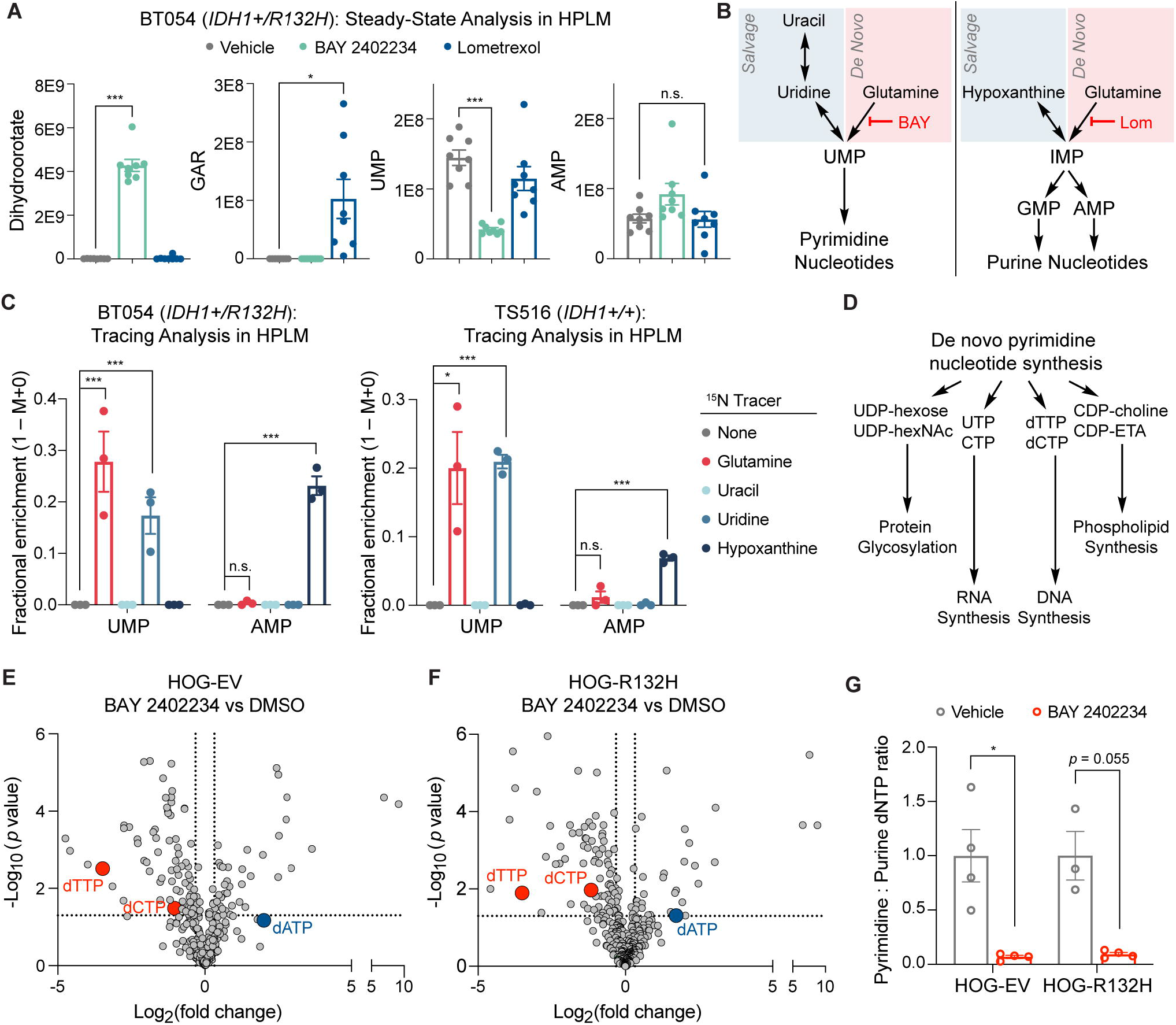
Glioma Cells Use Divergent Routes for Pyrimidine and Purine Nucleotide Synthesis Under Physiologic Conditions. (A) Steady-state quantification of the indicated metabolites by LC-MS in BT054 IDH1 mutant GSCs grown in human plasma-like medium (HPLM) and treated with 10 nM BAY 2402234, 5 μM lometrexol, or DMSO for 24 hours (*n =* 8 per condition). (B) Schema depicting pathways targeted by BAY 2402234 (BAY) and lometrexol (Lom). (C) ^15^N stable isotope tracing assays in BT054 (IDH1 mutant) and TS516 (IDH WT) GSC lines performed in HPLM (*n =* 3). ^15^N-labeled metabolites (listed at right) were administered to GSCs and labeling of representative pyrimidine [uridine monophosphate (UMP)] and purine [adenosine monophosphate (AMP)] nucleotides was measured 18 hours later by LC-MS. (D) Schema depicting select biological pathways utilizing intermediates derived from de novo pyrimidine nucleotide synthesis. (E and F) Volcano plots of metabolites measured by LC-MS in HOG-EV (E) or HOG-R132H (F) cells treated for 24 hours with 10 nM BAY 2402234 relative to DMSO (*n =* 4). (G) Ratio of pyrimidine (dTTP, dCTP) to purine (dATP) deoxynucleotide triphosphate (dNTP) pools in cells shown in (E) and (F). Note that our LC-MS method cannot distinguish dGTP from ATP; therefore, we report data for dATP only. For all panels, data presented are means ± SEM; \**p* < .05, \*\**p* < .01, \*\*\**p* < .001. n.s. = not significant. Two-tailed *p*-values were determined by unpaired *t*-test.

To test this idea, we conducted parallelized stable nitrogen isotope tracing assays for substrates that feed de novo and salvage nucleotide synthesis pathways and confirmed intracellular accumulation of nitrogen-labeled metabolites (Figure S4A). In support of our model, we found that the de novo pyrimidine synthesis pathway substrate ^15^N-glutamine labeled the majority of the UMP pool, with moderate and no labeling observed from the pyrimidine salvage pathway substrates ^15^N-uridine and ^15^N-uracil, respectively (Figure 7C). In contrast, the de novo purine synthesis pathway substrate ^15^N-glutamine failed to label AMP, while the purine salvage pathway substrate ^15^N-hypoxanthine robustly labeled AMP. Similar labeling patterns were observed in the IDH1/2 WT TS516 GSC line (Figure 7C). These data indicate that GSCs, independent of IDH status, engage qualitatively distinct mechanisms to produce pyrimidine and purine nucleotides, therefore partially explaining the differential impact of de novo pyrimidine and purine synthesis inhibitors in our drug screen.

To illuminate the mechanism by which *IDH1* oncogenes induce DHODH hyperdependence, we evaluated how DHODH inhibition affects the metabolomes of IDH1 mutant and IDH1 WT glioma cells. We focused on identifying changes in metabolic pathways that link pyrimidine synthesis with key pyrimidine-dependent cellular processes: protein glycosylation, RNA synthesis, phospholipid synthesis, and DNA synthesis (Figures 7D–F). Although BAY 2402234 treatment depleted some substrates for protein glycosylation (e.g. UDP-hexose) and phospholipid synthesis (e.g. CDP-choline), global levels of dolichol-linked oligosaccharides necessary for N-linked glycosylation and phosphatidylcholine and phosphatidylethanolamine lipids were largely unchanged (Figures S4B–E). BAY 2402234 also potently suppressed levels of the RNA precursors UTP and CTP (Figure S4F). However, upon reviewing data from our chemical library screen, we found that IDH1 mutant and IDH1 WT glioma cells were equally and exquisitely sensitive to treatment with the transcription inhibitor actinomycin D (Figure S4G). Therefore, the de novo pyrimidine synthesis vulnerability displayed by IDH1 mutant glioma cells is unlikely to reflect decreased transcription caused by decreased RNA precursors.

Our metabolomics study also revealed that DNA synthesis substrates were markedly dysregulated by DHODH inhibition. The pyrimidine deoxynucleotide triphosphates (dNTPs) dTTP and dCTP were potently suppressed while the purine dNTP dATP was upregulated (Figures 7E–G; Figure S5). Previous research revealed that imbalances in the ratio of pyrimidine to purine nucleotides required for DNA synthesis can evoke DNA damage (Kim et al., 2017), raising the possibility that IDH1 mutant cells are hypersensitive to the genotoxic effects of class-level perturbations to nucleotide abundance. In support of this idea, both engineered and patient-derived IDH1 mutant glioma cells displayed preferential induction of the DNA damage marker phospho-histone H2A.X (γH2A.X) in response to DHODH inhibition (Figures 8A and B). As an additional measure of DNA damage, we also characterized 53BP1 foci formation by immunofluorescence in isogenic IDH1 mutant and IDH1 WT HOG cells over a time course of BAY 2402234 treatment. 53BP1 foci were more abundant at baseline in IDH1 mutant versus IDH1 WT glioma cells (Figure 8C). This finding is consistent with an increase in 53BP1 foci observed in long-term hematopoietic stem cells exposed to chronic mutant IDH1 activity that was previously reported by a member of our group (T.W.M.) (Inoue et al., 2016). After 24–36 hours of DHODH inhibition, both IDH1 mutant and IDH1 WT cells exhibited increased numbers of 53BP1 foci per cell (Figure 8C). Interestingly, at 48 hours, 53BP1 foci increased further in IDH1 mutant cells, whereas 53BP1 foci number decreased markedly in IDH1 WT cells, thereby corroborating differential γH2A.X induction at this time point (compare Figure 8C to Figure 8A). These data suggest that persistent DNA damage accounts for the preferential cytotoxicity of de novo pyrimidine synthesis inhibitors against IDH1 mutant glioma cells versus IDH1 WT controls.

**Figure 8.**
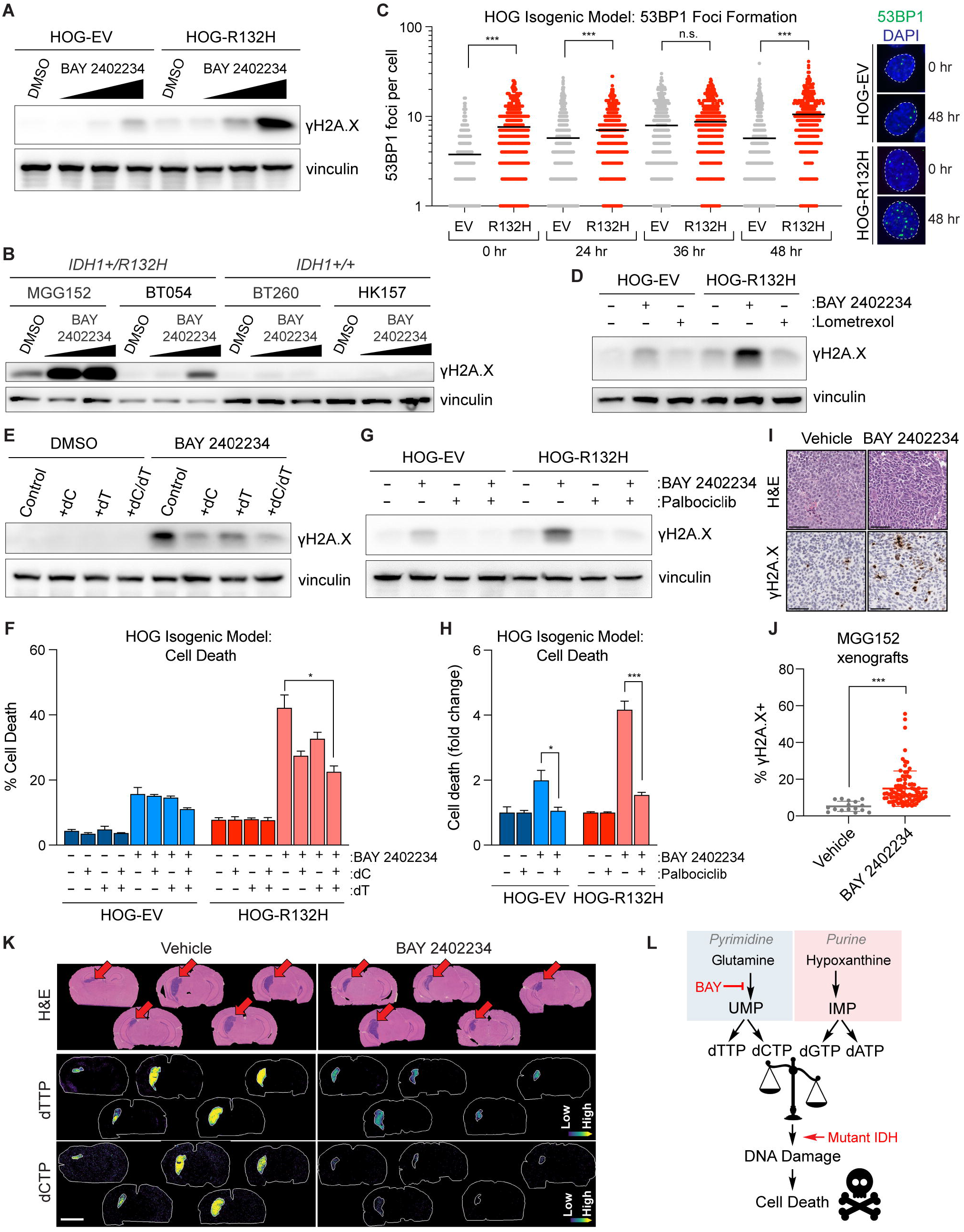
The *IDH1-R132H* Mutation Enhances DNA Damage Caused by DHODH Inhibition. (A) Immunoblot analysis of γH2A.X in HOG-EV and HOG-R132H cells treated with 1, 3, or 10 nM BAY 2402234 (indicated by triangles) or DMSO for 48 hours. (B) Immunoblot analysis of γH2A.X in IDH mutant (MGG152, BT054) and IDH wild-type (BT260, HK157) GSC lines treated with 3 or 10 nM BAY 2402234 (indicated by triangles) or DMSO. MGG152 cells were treated for 4 days, all others were treated for 5 days. (C) Quantification of 53BP1 foci formation in HOG-EV and HOG-R132H stable lines treated with 10 nM BAY 2402234 for the indicated durations (*n* ≥ 325 cells per condition per time point). Representative images from 0-and 48-hour time points are shown at right. (D) Immunoblot analysis of HOG-EV and HOG-R132H stable lines treated with 10 nM BAY 2402234, 5 μM lometrexol, or DMSO for 36 hours. (E and F) Immunoblot analysis of γH2A.X (E) and cell death assays (F) in HOG-EV and HOG-R132H stable lines treated with 10 nM BAY 2402234 or DMSO in the presence of 15 μM deoxycytidine (dC), 15 μM deoxythymidine (dT), both, or neither for 48 (E) or 72 hours (F) (*n =* 3). (G and H) Immunoblot analysis of γH2A.X (G) and cell death assays (H) in HOG-EV and HOG-R132H stable lines pre-treated with 1 μM palbociclib or DMSO for 48 hours, then treated with 10 nM BAY 2402234 or DMSO for 36 (G) or 72 (H) hours (*n =* 3). (I) Representative H&E and anti-γH2A.X IHC staining of MGG152 (IDH1 mutant) orthotopic glioma xenografts treated with BAY 2402234 (4 mg/kg PO QD) or vehicle for 3 days. Scale bar = 100 μm. (J) Quantification of γH2A.X-positive cells in MGG152 tumors treated as in (I). 16 regions of interest (ROIs) were analyzed from 2 vehicle-treated tumors and 86 ROIs were analyzed from 5 BAY 2402234-treated tumors. (K) Representative H&E-stained sections and MALDI-MSI ion images of brains from mice with MGG152 orthotopic xenografts treated as in (I) (*n* = 5 per cohort). Brains were harvested 4 hours after the final dose. Top panels: H&E staining showing site of tumor development. Middle and bottom panels: relative dTTP (middle) and dCTP (bottom) levels throughout the whole brain as determined by MALDI-MSI. Scale bars = 3 mm. (L) Schema depicting model by which BAY 2402234 (BAY) treatment preferentially kills IDH mutant glioma cells. For (C), data are means and variance is not displayed. For all other panels, data presented are means ± SEM. \**p* < .05, \*\**p* < .01, \*\*\**p* < .001, n.s. = not significant. Two-tailed *p*-values were determined by unpaired *t*-test.

To test the hypothesis that IDH1 mutant glioma cells are more susceptible to DNA damage-induced cell death caused by pyrimidine to purine dNTP pool imbalances, we first treated IDH1 mutant and IDH1 WT cells with lometrexol. As shown in Figures 7A and S1I, this de novo purine synthesis inhibitor does not affect purine nucleotide levels or fitness of glioma cells due to the predominance of the purine salvage pathway. In support of our hypothesis, we found that lometrexol treatment, unlike BAY 2402234, fails to induce DNA damage (Figure 8D). Furthermore, we found that rescuing pyrimidine dNTP pools by supplementing IDH1 mutant cells with deoxycytidine (dC) and deoxythymidine (dT) reduced γH2A.X and cell death upon DHODH inhibition (Figures 8E and F). Importantly, dC and dT provision during BAY 2402234 treatment did not restore pyrimidine substrates for protein glycosylation or phospholipid or RNA synthesis, suggesting that the mechanism of action does not principally involve these pathways (Figure S6A). Additionally, blocking cell cycle progression in G1-phase with the CDK4/6 inhibitor palbociclib attenuated DNA damage and cell killing by BAY 2402234 (Figures 8G and H), suggesting that suppression of the pyrimidine to purine dNTP ratio by BAY 2402234 elicits DNA damage that is dependent on replication stress during S-phase. Unlike palbociclib treatment and dC/dT supplementation, blocking 2HG synthesis with a mutant IDH1 inhibitor did not reverse DHODH inhibitor sensitivity in IDH1 mutant glioma cells (Figures S6B and C). Moreover, treating IDH1 WT cells with a cell permeable conjugate of (*R*)-2HG, (*R*)-2HG-TFMB, did not acutely sensitize them to DHODH inhibition (Figures S6D and E).

To extend our mechanistic studies in vivo, we quantified γH2A.X in BAY 2402234-or vehicle-treated orthotopic MGG152 IDH1 mutant glioma xenografts. In agreement with our in vitro studies (Figure 8B), we found that DHODH inhibition also increases DNA damage in vivo. Tumoral γH2A.X levels inversely correlated with the abundance of dTTP and dCTP (Figures 8I– K). Therefore, IDH1 mutant glioma cells sustain DNA damage in response to exhaustion of pyrimidine dNTP pools both in vitro and in vivo. Collectively, our data indicate that chronic mutant IDH activity sensitizes glioma cells to DHODH inhibition by increasing their susceptibility to replication-dependent DNA damage caused by nucleotide pool imbalances (Figure 8L).

## DISCUSSION

Our findings reveal that de novo pyrimidine nucleotide synthesis is a collateral vulnerability induced by *IDH* oncogenes in glioma. We used our recently developed MAPS screening platform to uncover drugs that preferentially reduce the fitness of IDH1 mutant glioma cells. This systematic, unbiased approach identified multiple classes of compounds that have previously been shown to exert enhanced antiproliferative activity against IDH mutant tumor cells. Our screen also identified three de novo pyrimidine synthesis antagonists: brequinar, 6-azauridine, and pyrazofurin. To translate this discovery, we first identified an alternative inhibitor of this pathway with high brain penetrance. We showed that BAY 2402234, a novel clinical-stage DHODH inhibitor, effectively targets DHODH in both normal and malignant brain tissues. Using a collection of engineered and patient-derived brain tumor models, we went on to show that BAY 2402234 monotherapy preferentially targets IDH1 mutant glioma cells both in vitro and in vivo. Notably, BAY 2402234 improved survival in a glioma mouse model that is resistant to IDH inhibitor treatment, highlighting this approach as a potential treatment strategy for patients who fail IDH inhibitor therapy. We also demonstrated efficacy of BAY 2402234 in models of lower grade (grade 3) gliomas, including a novel GEM model and organoid models, suggesting that BAY 2402234 is effective in both grade 3 and grade 4 tumors. The mechanism underpinning this activity involves both cell type-specific nucleotide biosynthesis pathway preference and mutant IDH1-driven susceptibility to DNA damage upon imbalance of pyrimidine and purine dNTP pools.

Our study complements prior investigations of alternative DHODH inhibitors, including teriflunomide, compound 10580, and brequinar, in IDH WT glioma xenograft model systems (Echizenya et al., 2020; Lafita-Navarro et al., 2020; Wang et al., 2019). Two groups have reported varying degrees of antitumor activity of DHODH inhibition in subcutaneous glioblastoma xenografts. Responses in this setting were linked to repression of protein glycosylation or rRNA synthesis. It is plausible that these DNA damage-independent effects of pyrimidine nucleotide depletion underpin the modest responses of IDH WT glioma cell lines to DHODH inhibition in vitro that we observed in our study. Contrasting these findings, both we and others observed minimal activity of DHODH inhibitor monotherapy against IDH WT glioblastoma xenografts when grown orthotopically (Wang et al., 2019). Similarly established IDH1 mutant xenografts, however, responded to BAY 2402234 treatment. Therefore, the tumor microenvironment, in addition to IDH mutational status, may play an important role in defining the response of brain tumors to this class of anticancer drugs. As DHODH inhibitors advance toward clinical testing in glioma, our work reveals that IDH1 mutations may serve as a targeted biomarker for trial enrollment. It is perhaps noteworthy in this regard that high DHODH expression is associated with inferior survival in patients with astrocytoma, IDH-mutant, grade 4 (Zhou et al., 2020).

We show that mutant IDH1 expression is sufficient to increase the susceptibility of glioma cells to DNA damage caused by imbalance of pyrimidine and purine dNTP pools. We observed that cell cycle progression is required for DNA damage to occur following pyrimidine nucleotide depletion, thereby implicating errors in DNA replication in this process. These findings build on extensive prior research establishing nucleotide pool homeostasis as a critical determinant of DNA replication fidelity (Lee et al., 2018; Meuth, 1989). Moreover, our work extends the idea that certain tumor cell oncogenotypes increase pyrimidine synthesis dependence and sensitivity to nucleotide pool imbalance. Lung cancer cells with mutations in *KRAS* and *LKB1* engage a non-canonical pyrimidine biosynthesis program dependent on carbamoyl phosphate synthetase-1 (CPS1) to achieve pyrimidine and purine nucleotide balance and avert DNA damage (Kim et al., 2017). Observing similar vulnerabilities to nucleotide imbalance driven by *IDH1* and *KRAS*/*LKB1* mutations in brain and lung tumors, respectively, suggests that this liability could be associated with other oncogenes and tumor subtypes, thus representing a possible paradigm that could be therapeutically exploited for a host of tumors.

*IDH* mutations have been previously linked with DNA damage susceptibility in glioma, in part through repression of homology-directed repair (HDR) (Sulkowski et al., 2017). (*R*)-2HG inhibits the KDM4 family of histone demethylases that act on methylated histone 3 lysine 9 (H3K9) residues. Global hypermethylation of H3K9 residues in IDH mutant tumor cells can impede recognition of H3K9me3 marks specifically deposited at sites of DNA double strand breaks by proteins involved in HDR (Sulkowski et al., 2020). However, this process cannot fully explain the increased DNA damage sensitivity in our models because H3K9 hypermethylation and HDR suppression are readily reversed by mutant IDH1 inhibition and (*R*)-2HG depletion. In contrast, sensitivity of IDH1 mutant glioma cells to DHODH inhibitors, although correlating with steady-state (*R*)-2HG levels (see Figures 1L and 2H), is not acutely altered by abrupt changes in (*R*)-2HG levels. In this regard, Lu and colleagues showed that engineering IDH1 mutations in U251 glioma cells increases their sensitivity to the DNA damaging agent temozolomide via a mechanism that is likewise insensitive to short-term mutant IDH1 inhibitor treatment (Lu et al., 2017). IDH1 mutations are known to cause both reversible and irreversible changes in chromatin structure and gene expression (Turcan et al., 2018), and irreversible changes may underpin durable DNA damage hypersensitivity phenotypes in IDH1 mutant glioma cells. In the context of DHODH inhibition, DNA damage may also be reinforced by decreased flux from glucose to pyrimidine nucleotides, which has been previously observed in IDH1 mutant glioma cells (Garrett et al., 2018).

In addition to nominating a new therapeutic target in IDH1 mutant glioma, we also report the development of a novel genetically engineered mouse model of this disease. By introducing clinically relevant mutations that are frequently found in astrocytoma, IDH-mutant, grade 3 into neural cells of adult mice, we generated autochthonous gliomas that bear striking genetic and histologic resemblance to these human tumors. Notably, both our model and a previously reported GEM model of astrocytoma, IDH-mutant, grade 4 developed by Philip and colleagues (Philip et al., 2018) demonstrate that the *IDH1-R132H* oncogene drives gliomagenesis when expressed in the presence of naturally co-occurring mutations. These advances circumvent longstanding challenges in developing genetically engineered models of glioma that depend on the oncogenic functions of *IDH* mutations. *IDH1-R132H* expression alone is insufficient to cause glioma formation in mice (Bardella et al., 2016; Sasaki et al., 2012a) and can impede gliomagenesis when coupled with mutations that are infrequently observed in IDH1 mutant human brain tumors (Núñez et al., 2019). We show that the *IDH1-R132H* oncogene cooperates with mutations that activate PI3K signaling to promote neural cell transformation. This finding corroborates and underscores the significance of a previously identified association between *PIK3R1* mutations and poor outcomes in patients with IDH mutant astrocytoma (Aoki et al., 2018). Our GEM model provides an in vivo platform to study oncogenic mechanisms and treatment response profiles of mutant IDH1-driven glioma and opens new avenues to explore mutant IDH1 action in the setting of treatment-naïve, lower-grade brain tumors, which have been difficult to model effectively to date.

Although the discovery of IDH mutations spurred optimism that inhibitors of IDH mutant oncoproteins might display broad activity against IDH mutant gliomas, these agents have not shown benefit in patients with contrast-enhancing brain tumors (Mellinghoff et al., 2021), with a 0% objective response rate reported in a recent phase I trial. De novo resistance to mutant IDH inhibitors may be explained by the observation that mutant IDH oncogenes elicit durable changes in gene expression that are insensitive to acute (*R*)-2HG depletion, thus transforming neural cells through a “hit-and-run” mechanism (Johannessen et al., 2016). Therefore, identifying therapies that are effective against IDH mutant gliomas independently of whether they display an ongoing requirement for (*R*)-2HG is a key challenge in the field. Our data show that two mouse models of IDH1 mutant glioma (one grade 3 model and one grade 4 model) respond to DHODH inhibitor monotherapy. Importantly, the grade 4 model has previously been shown by Tateishi and colleagues to be refractory to mutant IDH1 inhibition and (*R*)-2HG depletion (Tateishi et al., 2015). Our work provides preclinical rationale to initiate clinical studies of BAY 2402234 for brain tumor therapy by identifying IDH mutational status as a predictive biomarker of response to this agent. Furthermore, we outline a new therapeutic strategy that shows promise for treating IDH1 mutant gliomas that display de novo resistance to mutant IDH1 inhibitors, thereby directly addressing an important yet unmet clinical need.

## ACKNOWLEDGEMENTS

We thank C. Stiles (DFCI), T. Batchelor (BWH), and S. Morrison (UTSW) for their insightful feedback on our work; J. Alberta (DFCI) for sharing Olig2 antibody; J. Brugge (HMS) for supporting development of the MAPS platform; Kaelin, McBrayer, Abdullah, DeBerardinis, and Losman laboratory members for helpful discussions; I. Mellinghoff (MSKCC), S. Weiss (Univ. of Calgary), and R. Pieper (UCSF) for sharing cell lines; Bayer Pharmaceuticals for sharing BAY 2402234; M. Yuan (Harvard) for help with mass spectrometry; and DFCI Animal Resources Facility and UTSW Animal Resource Center for help with mouse work. This study was supported by awards from the Broad-Bayer Alliance (Joint Project Proposal Grant 7000062-5500001463) and from NIH/NCI: K22CA237752 to S.K.M., R01CA258586 to S.K.M. and K.G.A., and P50CA165962 to N.Y.R.A, D.P.C., K.L.L., and W.G.K. (and related Career Enhancement Project award to S.K.M.), P50CA211015 to H.I.K., U54CA210180 to N.Y.R.A, P41EB028741 to N.Y.R.A, and T32EB025823 to S.A.S. N.Y.R.A. is supported by the Pediatric Low-Grade Astrocytoma Program at PBTF. Drug screening using MAPS by J.E.E. and I.S.H was supported by the Ludwig Center at Harvard. Organoid modeling work was partly supported by an award from Oligo Nation Foundation to S.K.M. and K.G.A. S.K.M. is supported by a Cancer Prevention and Research Institute of Texas (CPRIT) award (RR190034), a V Scholar Award from the V Foundation for Cancer Research (V2020-006), a Distinguished Scientist Award from the Sontag Foundation, and a gift from the Jonesville Foundation. D.D.S. was supported by the HHMI Medical Research Fellows Program and the Scholars in Medicine Program at Harvard Medical School. K.G.A. is supported by the Eugene P. Frenkel, M.D. Endowment. C.E.B. is supported by an award from the Burroughs Wellcome Trust.

## AUTHOR CONTRIBUTIONS

D.D.S. and S.K.M. did the experiments and, together with K.G.A. and W.G.K., designed the experiments, analyzed data, and wrote the manuscript. M.R.S. and M.M.L. did stable isotope tracing and metabolomics experiments. A.C.W., W.G., and J.K. helped create the astrocytoma GEM model. J.E.E. and I.S.H. developed and used the MAPS compound screening platform. C.E.B. and J.B. established and treated organoid models and L.C.G. managed related clinical data. S.A.S. and M.S.R. did MALDI-MSI experiments and interpreted the data with N.Y.R.A. Y.-F.L. and M.X. performed DNA damage assays and analyzed data with P.L. B.H. and S.L. helped genetically engineer NHA cells and evaluated their transformation status. R.B.J. did immunohistochemistry and analyzed results with S.S. D.B. performed MRI studies and interpreted data with Q.-D.N. M.S.M.-S., J.M.A., L.G.Z., and T.P.M. conducted and/or analyzed LC-MS studies. R.E.L. synthesized (*R*)-2HG-TFMB. T.D. quantified N-linked glycans and interpreted data with M.A.L. H.I.K., T.W.M., D.P.C., and K.L.L. provided GSC and/or mouse models and guidance on their use. S.G., A.S., M.J., and A.J. provided BAY 2402234 and guidance on its use in preclinical models. K.L.L. and T.E.R. performed neuropathology evaluations of tissues from our GEM model and organoid models, respectively. D.P.C., R.J.D., and K.L.L. helped design and interpret experiments.

## DECLARATION OF INTERESTS

R.J.D., W.G.K., and S.K.M. have served as paid advisors to Agios Pharmaceuticals. W.G.K. receives compensation for his roles as Eli Lilly and Company Board Director and founder of Tango Therapeutics and Cedilla Therapeutics. K.L.L. receives research support from Eli Lilly and Company via the DFCI. S.K.M. and W.G.K. received research funding from Bayer Pharmaceuticals. Bayer had no influence over the design, execution, or interpretation of studies. N.Y.R.A. is key opinion leader for Bruker Daltonics, scientific advisor to Invicro, and receives support from Thermo Finnegan and EMD Serono. D.P.C. has consulted for Lilly, GlaxoSmithKline, Boston Pharmaceuticals and serves on the advisory board of Pyramid Biosciences, which includes an equity interest. All other authors declare no competing interests.

## MATERIALS AND METHODS

### Cell lines

HOG cells (human oligodendroglioma line from a male) were a gift of P. Paez, SUNY Univ. at Buffalo and were cultured in IMDM medium with 10% FBS and 1% penicillin/streptomycin. Stable HOG cell lines expressing either EV or IDH1-R132H were generated using plenti-Ubc-IRES-hygro plasmid backbone and cDNA cloning strategies as previously described (McBrayer et al., 2018). Stable cell lines were selected with 500 µg/mL hygromycin and maintained in media the above media supplemented with 200 µg/mL hygromycin and were cultured for at least 5 weeks after selection prior to experimentation. NHA cells (human astrocytes immortalized with HPV E6 and E7 and hTERT) (Sonoda et al., 2001) (sex unknown) were a kind gift of Dr. Russell Pieper (UCSF). NHA cells were cultured in DMEM medium (Gibco 11995-065) containing 10% FBS and 1% penicillin/streptomycin. Stable, late passage NHA cells expressing either EV or IDH1-R132H used in this study were described previously (Koivunen et al., 2012) and maintained in media supplemented with 500 µg/mL hygromycin. ***I****dh1^LSL-R132H/+^*;*LSL-**C**as9^+/-^* (***IC***) MEFs created in this study were cultured in DMEM medium (Gibco 11995-065) containing 10% FBS and 1% penicillin/streptomycin.

All HOG, NHA, and MEFs lines were cultured in the presence of 5% CO_2_ and ambient oxygen at 37 °C. All cell lines were routinely evaluated for mycoplasma contamination and tested negative throughout the study. Cell line authentication was not performed because reference short term tandem repeat profiles have not been established for these cell lines.

### Primary Cell Cultures

GSC lines TS516 and TS603 (sexes unknown) were obtained from I. Mellinghoff at MSKCC (Rohle et al., 2013). BT054 cells (female) were obtained from S. Weiss at Univ. of Calgary (Kelly et al., 2010). BT260 cells (sex unknown) were obtained from K. Ligon at DFCI (Koivunen et al., 2012). HK211 (female), HK213 (male), HK252 (male), HK308 (female), and HK157 (female) cells were obtained from H. Kornblum at UCLA (Laks et al., 2016). TS516, TS603, BT054, and BT260 cells were cultured in NeuroCult NS-A Basal Medium (Human) with Proliferation Supplement (StemCell Technologies 05750) supplemented with EGF (20 ng/mL), bFGF (20 ng/mL), heparin (2 μg/mL), 1% penicillin/streptomycin, amphotericin B (250 ng/mL), and Plasmocin (2.5 μg/mL). HK213, HK211, HK308, and HK157 cells were cultured in DMEM-F12 medium (Gibco 11320033) supplemented with 3 mM glutamine, 1x B27, EGF (20 ng/mL), bFGF (20 ng/mL), heparin (2 μg/mL), 0.5% penicillin/streptomycin, amphotericin B (125 ng/mL), and Plasmocin (2.5 μg/mL). MGG152 cells (Wakimoto et al., 2014) were cultured in Neurobasal Medium (Gibco 21103049) supplemented with 3 mM glutamine, 1x B27, 0.25x N2, EGF (20 ng/mL), bFGF (20 ng/mL), heparin (2 μg/mL), 0.5% penicillin/streptomycin, amphotericin B (125 ng/mL), and Plasmocin (2.5 μg/mL). DF-AA27 GEMM GSCs were cultured in NeuroCult Basal Medium (Mouse and Rat) with Proliferation Supplement (StemCell Technologies 05702) supplemented with EGF (20 ng/mL), bFGF (20 ng/mL), heparin (2 μg/mL), 1% penicillin/streptomycin, amphotericin B (250 ng/mL), and Plasmocin (2.5 μg/mL). Primary human astrocytes (sex unknown) were purchased from Lonza (CC-2565) and cultured in ABM Basal Medium (Lonza CC-3187) supplemented with AGM SingleQuots Supplements (Lonza CC-4123). Human neural stem cells (sex unknown) were purchased from Fisher Scientific (10419428) and cultured in DMEM/F12 medium supplemented with 3 mM glutamine, EGF (20 ng/mL), bFGF (20 ng/mL), heparin (2 μg/mL), 0.5% penicillin/streptomycin, amphotericin B (125 ng/mL), Plasmocin (2.5 μg/mL), and StemPro Neural Supplement (Thermo Fisher A1050801). All GSC lines and human neural stem cells were maintained as neurospheres using ultra low-adherence culture dishes and dissociated 1-2 times per week with Accutase (StemCell Technologies 07922). Primary human astrocytes were cultured adherently on plates coated with Geltrex (Life Technologies A1413202). Sex and source of each line is stated above and listed as unknown if unreported in the original publication describing its derivation.

All human GSC lines were cultured in the presence of 5% CO_2_ and ambient oxygen at 37 °C. DF-AA27 mouse GSCs were cultured in the presence of 5% CO_2_, 5% O_2_ at 37 °C. All cell lines were routinely evaluated for mycoplasma contamination and tested negative throughout the study. Cell line authentication was not performed because reference short term tandem repeat profiles have not been established for these cell lines. Only low passage GSC lines were used and these cells were discarded after 3 months in culture to prevent genetic and/or phenotypic drift.

### Animals

All care and treatment of experimental animals were carried out in strict accordance with Good Animal Practice as defined by the US Office of Laboratory Animal Welfare and approved by the Dana-Farber Cancer Institute (protocol 04-019) or the UT Southwestern Medical Center (protocol 2019-102795) Institutional Animal Care and Use Committee. Animal welfare assessments were carried out daily during treatment periods. Female mice were housed together (2–5 mice per cage) and provided free access to standard diet and water. Mice were randomized to experimental arms prior to cell implantation and/or treatment. For orthotopic glioma cell implantations and intracranial AAV injections, mice were anesthetized via intraperitoneal injection of ketamine (140 mg/kg) and xylazine (12 mg/kg) and immobilized using a stereotactic frame. An incision was made to expose the skull surface, and a hole was drilled into the skull. AAV (1 μL) or cells suspended in 1–3 μL of 2% FBS in PBS (NHA cells) or cell culture medium (GSC lines) were injected into the brain through the hole using a 5 μL syringe (Hamilton). The skin was closed with surgical clips, and buprenorphine was given for analgesia. Tumor size and survival analyses were performed by researchers who were not blinded to the treatment arms or genotypes of the mice. Mice were euthanized when they either displayed neurological symptoms or became moribund.

#### Orthotopic xenograft and allograft models

NHA orthotopic xenografts were created by intracranial injection of 3 × 10^5^ cells expressing either IDH1-R132H, PIK3R1-D560_S565del, both, or neither or H-Ras-V12 into female NCr nude mice (Taconic). Cells were implanted 1 mm anterior and 2 mm lateral to the lambda, 2.5 mm below the surface of the brain. All injected cells expressed a firefly luciferase-IRES-GFP bicistronic expression cassette (LeGO-iG2-FLuc vector) as previously described (McBrayer et al., 2018) to allow for non-invasive bioluminescence imaging of tumor growth. Mice were imaged once per week following cell injection. In vivo imaging was performed after intraperitoneal injection of luciferin (50 mg/kg). Mice were anesthetized with isoflurane and imaged using an IVIS camera (PerkinElmer). Imaging data were analyzed using Living Image software (PerkinElmer). Tumor growth rates were determined by subtracting the initial signal from the signal at the imaging time point immediately preceding euthanasia or at 16 weeks after cell implantation (for mice that did not require euthanasia during study period) and dividing by the number of intervening weeks.

MGG152 and TS516 orthotopic xenografts were created by intracranial injection of 1 × 10^5^ cells into female ICR SCID mice (Taconic). MGG152 cells were implanted 2 mm lateral to the bregma, 2.5 mm below the surface of the brain. TS516 cells were implanted 1 mm posterior and 2 mm lateral to the bregma, 2 mm below the surface of the brain. For survival studies, MGG152 and TS516 tumor-bearing mice were randomized to treatment arms 12 and 9 days after tumor cell implantation, respectively. Immediately following randomization, BAY 2402234 (4 mg/kg PO QD) or vehicle treatments were dosed continuously until mice were euthanized. Radiation (9 Gy in 3 fractions QOD) or sham treatments started two days after randomization. Mice undergoing radiation or sham treatments were anesthetized via intraperitoneal injection of ketamine/xylazine and placed in a lead shield covering their bodies but not their heads. Radiation was delivered using a Gammacell-40 irradiator (Nordion). For MALDI-MSI studies, brains were harvested from MGG152 tumor-bearing mice 33 days after tumor cell implantation.

DF-AA27 orthotopic allografts were created by intracranial injection of 1 × 10^5^ cells into female ICR SCID mice (Taconic). DF-AA27 cells were implanted 1 mm anterior and 2 mm lateral to the lambda, 2.5 mm below the surface of the brain. For tumor volume measurement studies, DF-AA27 tumor-bearing mice were randomized to BAY 2402234 or vehicle treatment arms upon tumor detection by MRI. Treatment commenced immediately following randomization and MRIs were acquired again 3 weeks later.

#### Genetically engineered mouse models

Intracranial injection of AAV was performed using 1.5-6 month old male and female transgenic mice (genotypes indicated in Figure 5A). Strains used were: H11^LSL-Cas9^ (B6;129-*Igs2^tm1(CAG-cas9*)Mmw^*/J, Jackson stock number 026816) (Chiou et al., 2015), R26-Pik3ca^H1047R^ (FVB.129S6-*Gt(ROSA)26Sor^tm1(Pik3ca*H1047R)Egan^*/J, Jackson stock number 016977) (Adams et al., 2011), Idh1^tm1Mak^ (backcrossed to C57BL/6, provided by T. Mak at University of Toronto) (Sasaki et al., 2012a, 2012b). To generate the mouse strains used for the injection scheme outlined in Figure 5A, we first bred transgenic H11^LSL-Cas9+/+^ mice with Idh1^tm1Mak/+^ mice to produce H11^LSL-Cas9+/-^;Idh1^tm1Mak/WT^ progeny, which were subsequently interbred to produce H11^LSL-Cas9+/+^;Idh1^tm1Mak/WT^ mice. H11^LSL-Cas9+/+^;Idh1^tm1Mak/WT^ mice were crossed with R26-Pik3ca^H1047R+/+^ to produce ***PIC*** (H11^LSL-Cas9+/-^;Idh1^tm1Mak/WT^;R26-Pik3ca^H1047R+/-^) and ***PC*** (H11^LSL-Cas9+/-^;Idh1^WT/WT^;R26-Pik3ca^H1047R+/-^) mice. Separately, H11^LSL-Cas9+/+^;Idh1^tm1Mak/WT^ mice were crossed with wild-type FVB mice to produce ***IC*** (H11^LSL-Cas9+/-^;Idh1^tm1Mak/WT^) and ***C*** (H11^LSL-Cas9+/-^;Idh1^WT/WT^) mice. Genotyping was performed by Transnetyx. For intracranial AAV injections, mice were first anesthetized via intraperitoneal injection of ketamine (140 mg/kg) and xylazine (12 mg/kg). Mice were immobilized using a stereotactic frame. Following skin incision to expose the skull, a hole was drilled 1 mm posterior and 1 mm lateral to the bregma. 1 μL of virus at 1 × 10^13^ to 1 × 10^14^ virions/mL, (1 × 10^10^ to 1 × 10^11^ total AAV particles) was injected 2.1 mm below the surface of the brain over 5 minutes. Mice were monitored with serial monthly MRI scans, and glioma incidence was determined by histopathologic evaluation of brain tissues after euthanasia and/or intracranial tumor detection by MRI.

### Human Subjects

The study was conducted according to the principles of the Declaration of Helsinki. Patient tissue and blood were collected following ethical and technical guidelines on the use of human samples for biomedical research at UT Southwestern Medical Center after informed patient consent under a protocol approved by UT Southwestern Medical Center’s Institutional Review Board. Use of human brain tissue was organized by the Department of Clinical Pathology at UT Southwestern Medical Center. All patient samples were de-identified before processing. All patient samples and organoids were diagnosed and graded according to the 2021 *WHO Classification of Tumours of the Central Nervous System (CNS)*, 5^th^ edition (Louis et al., 2021). Organoid models were created from patients with the following ages and sexes: UTSW6577 (32 male), UTSW1382 (27 male), UTSW2243 (53 female), UTSW9647 (33 male), UTSW2846 (53 male), UTSW2294 (59 male), UTSW9698 (66 female), UTSW9501 (76 male).

Organoid creation from primary glioma tissue was conducted as described previously (Abdullah et al., 2021). Briefly, tumor tissue was collected from the operating room and suspended in ice cold Hibernate A (BrainBits HA). Tumor pieces were exposed to RBC lysis buffer (Thermo Fisher 00433357) and washed with Hibernate A containing Glutamax (final conc. = 2 mM, Thermo Fisher 35050061), penicillin/streptomycin (final conc. = 100 U/mL and 100 μg/mL, respectively, Thermo Fisher 15140122), and Amphotericin B (final conc. = 0.25 μg/mL, Gemini Bio-Products 400104). Tissues were cut using dissection scissors into 1-2 mm^3^ pieces and suspended in 1 mL Short-Term Glioma Organoid Medium (formulation provided below). One organoid in 1 mL Short-Term Glioma Organoid Medium was plated per well of a 24-well ultra-low adherence plate. Plates were rotated at 120 rpm in a CO_2_-resistant shaker (Fisher Scientific 88-881-103) in a humidified incubator at 37°C, 5% CO_2_, and 5% O_2_. Short-Term Glioma Organoid Medium was refreshed in organoid cultures every 48 hours. All organoids were cultured at least four weeks before treatment. Organoids were randomized to BAY 2402234 or vehicle treatments and treated for 2 weeks before being fixed for IHC analysis. Formulation of Short-Term Glioma Organoid Medium is as follows: 48 mL Long-Term Glioma Organoid Medium, 48 µl 2-mercaptoethanol (BME) (final conc. = 55 µM, Thermo Fisher BP176-100), and 12 μl human insulin (final conc. = 2.375-2.875 μg/mL, Sigma Aldrich I9278). Formulation of Long-Term Glioma Organoid Medium is as follows: 250 mL DMEM:F12 medium (Thermo Fisher 1132033), 250 mL Neurobasal medium (Thermo Fisher 21103049), 5 mL 100X Glutamax (final conc. = 2 mM), 5 mL 100X low-glutamate non-essential amino acids mixture (final conc.: Gly, L-Ala, L-Asn, L-Asp, L-Pro, L-Ser = 100 μM and L-Glu = 300 nM), 5 mL penicillin/streptomycin (final conc. = 100 U/mL and 100 μg/mL, respectively), 10 mL B-27 Supplement without Vitamin A (Thermo Fisher 12587010), 5 mL N-2 Supplement (Thermo Fisher 17502048). Long-Term Glioma Organoid Medium and Short-Term Glioma Organoid Medium stocks were used up to 2 months and 1 week after preparation, respectively.

## METHOD DETAILS

### Chemicals

TFMB ester of (*R*)-2-hydroxyglutarate was generated as previously described by R. Looper at University of Utah (Losman et al., 2013). BAY 2402234 was provided by Bayer Pharmaceuticals. Where indicated, cell culture media also contained the following additives: brequinar (Sigma-Aldrich), 6-azauridine (Sigma-Aldrich), pyrazofurin (Sigma-Aldrich), uridine (Sigma-Aldrich), lometrexol (Cayman), AGI-5198 (XcessBio), deoxythymidine (Sigma-Aldrich), deoxycytidine (Sigma-Aldrich), palbociclib (Cayman), tunicamycin (Sigma-Aldrich).

### Vectors

Empty vector (plenti-Ubc-IRES-hygro) and IDH1-R132H (plenti-Ubc-IDH1-R132H-HA-IRES-hygro) lentiviral expression plasmids used to create stable HOG cell lines were previously described (McBrayer et al., 2018). Empty vector (pBABE-HA-hygro) and IDH1-R132H (pBABE-IDH1-R132H-HA-hygro) retroviral expression plasmids used to create stable NHA cell lines were previously described (Koivunen et al., 2012). PIK3R1 expression vectors (WT, D560_S565del, R574fs, T576del) used to generate NHA stable cell lines were from Addgene (pLenti-Flag-P85, Addgene 40219; pLenti4-Flag-P85-DKRMNS560del, Addgene 40225; pLenti-Flag-P85-R574fs, Addgene 40227; pLenti4-Flag-P85-T576del, Addgene 40228) (Quayle et al., 2012). H-Ras-V12 expression vector was from Addgene (pLXSN-H-Ras_V12, Addgene 39516). The lentiviral plasmid used to express GFP and firefly luciferase markers in NHA stable cells, LeGO-iG2-FLuc, was previously described (McBrayer et al., 2018). Retroviral vectors used to immortalize (pLBCX-Large-T-antigen^K1 mutant^) and genetically engineer (pMSCV-MerCreMer-hygro) ***I****dh1^LSL-R132H/+^*;*LSL-**C**as9^+/-^* (***IC***) MEFs were previously generated in the Kaelin laboratory. To make pLBCX-Large-T-antigen^K1 mutant^, Large-T-antigen^K1 mutant^ cDNA was PCR amplified using a 5’ primer that introduced a SalI site and a 3’ primer that introduced a ClaI site. The PCR product and pLBCX empty vector were digested with SalI and ClaI, gel-purified, and ligated. To make pMSCV-MerCreMer-hygro, MerCreMer cDNA was PCR amplified using a 5’ primer that introduced a NotI site and a 3’ primer that introduced a PacI site. The PCR product and pMSCV-hygro empty vector were digested with NotI and PacI, gel-purified, and ligated.

The pAAV2-sgTrp53-sgAtrx-EFS-Cre AAV vector was created using an approach similar to that described previously (Oser et al., 2019). Effective sgRNAs (sequences in Key Resources Table) targeting mouse *Trp53* and *Atrx* genes were identified from a previous publication (Platt et al., 2014) and empirically, respectively. To generate the AAV vector depicted in Figure 5A, we first generated a destination vector (pAAVGao-DEST-EFS-Cre-spA). This was accomplished by inserting a multiple cloning site sequence between XbaI and NotI sites in pX551 (a gift from F. Zhang at MIT, Addgene 60957). We then inserted the universal gateway cassette between PacI and NheI using restriction enzymes to generate an intermediate vector named pAAVGao-DEST. Next, we performed overlapping PCR to assemble the EFS promoter, Cre, and short poly(A) signal that contained a 5′ PacI site and 3′ XbaI site. The PCR product was digested with PacI and XbaI and ligated into pAAVGao-DEST cut with these two enzymes. Next, we performed Gibson assembly to generate an entry vector (pENTR223-sgTrp53-sgAtrx) containing sgRNAs targeting *Trp53* and *Atrx*. pENTR223-sgTrp53-sgAtrx was then mixed with pAAVGao-DEST-EFS-Cre-spA to make the final pAAV2-sgTrp53-sgAtrx-EFS-Cre by homologous recombination reaction using LR Clonase II (Life Technologies 11791100) at 25°C for 1 h per the manufacturer’s instructions. The reaction mixtures were then transformed at a ratio of 1:10 (volume recombination reaction:volume competent cells) into HB101 cells and ampicillin-resistant colonies were screened by restriction digestion of miniprep DNA and subsequently validated by whole plasmid DNA sequencing at the MGH CCIB DNA Core. AAV packaging and titering using pAAV2-sgTrp53-sgAtrx-EFS-Cre was performed by Vigene Biosciences.

### Transient transfection and in vitro viral transduction

Lentiviral and retroviral particles were made by Lipofectamine 2000-based cotransfection of HEK293T cells with expression vectors and packaging plasmids psPAX2 (Addgene 12260) and pMD2.G (Addgene 12259) or gag/pol (Addgene 14887) and VSV.G (Addgene 14888), respectively, in a ratio of 4:3:1. Virus-containing media was collected 48 and 72 hours after transfection, passed through a 0.45 μm filter, divided into 1 ml aliquots, and frozen at −80°C until use.

NHA cells or MEFs were plated at a density of 0.3 × 106 cells per well in a 6-well plate. The next day, 2.4 μL polybrene in 0.5 mL media was added to each well in addition to 0.5 mL viral supernatant. Plates were centrifuged at 4,000 × g for 30 minutes at room temperature and incubated overnight. The following day, cells were expanded and replated. Stable cell lines were selected in 500 μg/mL hygromycin, 2 μg/mL puromycin, 1 mg/mL G418, or 10 μg/mL blasticidin based on the drug selection cassette present in each vector of interest.

### Creating genetically engineered MEFs

A dam (*LSL-Cas9^+/+^*), impregnated by a male (*Idh1^LSL-R132H/+^*) during a timed mating, was euthanized for embryo harvest at day E13.5. Fibroblasts were isolated from an embryo later determined to be heterozygous for both *Cas9* and mutant *Idh1* alleles (***IC*** genotype). MEFs were immortalized by transduction with retroviral particles produced from the pLBCX-Large-T-antigen^K1 mutant^ vector. Next, immortalized MEFs were transduced with retroviral particles produced from the pMSCV-MerCreMer-hygro vector to constitutively express tamoxifen-inducible Cre recombinase (Sohal et al., 2001) or were mock infected. After selection, MerCreMer-expressing MEFs were treated with 1 μM 4-hydroxytamoxifen for 6 days to activate Cre.

### DF-AA27 GSC line creation

After observing brain tumor formation in an AAV-injected ***PIC*** mouse by MRI, the mouse was euthanized and brain tissue was harvested. A portion of the right hemisphere of the brain containing tumor tissue was isolated and dissociated to a single cell suspension using the Neural Tissue Dissociation Kit (Miltenyi) and a gentleMACS Dissociator instrument (Miltenyi). Cells were cultured in 5% CO_2_ and 5% O_2_ on ultra-low adherence plates to select for neurosphere-forming cells. The resulting murine GSC line was named DF-AA27. Notably, prior attempts to generate murine GSC lines from this GEM model under ambient oxygen conditions were not successful.

### Drug Screen

The MAPS platform (Harris et al., 2019) was used to test both a commercial anticancer drug library and a custom-curated metabolic inhibitor drug library. Screen was performed at the ICCB-Longwood Screening Facility (https://iccb.med.harvard.edu/small-molecule-screening). HOG-EV or HOG-R132H cells were seeded at a density of 500 cells per well in a final volume of 30 μL per well of 384-well plates. After 24 hours, a Seiko Compound Transfer Robot pin transferred 100 nL of each drug library into wells with plated cells. Following pin-transfer, 20 μL of cell culture medium was added to all wells, resulting in each drug being applied at a final 10-point concentration series ranging from 20 μM to 1 nM. After 72 hours of drug treatment, the cells were washed with PBS, fixed with 4% formaldehyde, and stained with 5 mg/mL bisbenzimide. An Acumen Cellista plate cytometer was used to image plates and determine the cell numbers in individual wells. XY plots were generated comparing relative numbers of surviving HOG-EV and HOG-R132H cells with concentrations of each drug tested. Area under the curve (AUC) values were calculated for each plot and drugs were ranked based on the difference between the AUCs for HOG-EV and HOG-R132H cells. Hits were defined as drugs that displayed >10 %AUC_Diff_ values, where %AUC_Diff_ = ((AUC_EV_ - AUC_R132H_)/AUC_EV_)*100. Using these criteria, 56 hits were identified, including the 36 highest ranking hits displayed in Figure 1B.

### Cell death quantification

Cells were stained with AnnexinV-FITC (BD Biosciences 556547) according to manufacturer’s instructions and DAPI (100 ng/mL final concentration) to identify early and late apoptotic cells, respectively. Cells were analyzed using either an LSR II (BD Biosciences) or LSR Fortessa (BD Biosciences) flow cytometer and data was processed using FCS Express software (De Novo). Dead cells included those that were AnnexinV-FITC^+^/DAPI^-^, AnnexinV-FITC^-^/DAPI^+^, or AnnexinV-FITC^+^/DAPI^+^.

### Doubling time calculation

Doubling times of human GSC lines were calculated by plating 2 x 10^5^ cells per well in a 6-well plate. 5 days later, cells were counted using a Vi-Cell XR (Beckman Coulter) cell viability analyzer. Doubling time (T_d_) was calculated using the following formula:

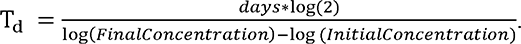

### In vitro pyrimidine synthesis inhibition

Pyrimidine synthesis inhibitor treatments of HOG cells were performed by plating HOG cells at a density of 5,000-10,000 cells per cm^2^ with or without 100 μM uridine supplementation. 24 hours later, the drug being tested was added at the indicated concentrations, and cell death was analyzed at the indicated time points. For GSC lines, cells were plated in 24-well ultra-low-attachment plates at a density of 50,000 cells per well with 0.25 mL media per well. An additional 0.25 mL medium containing 2x the desired drug concentrations was added at the time of plating to achieve the final concentrations of drug in 0.5 mL media per well. Fresh media (0.5 mL) with drug or vehicle was added every 4 days. In Figures 1 and 2, human GSCs were harvested for cell death quantification after two population doublings (5 days for TS516 and HK308; 6 days for BT260; 7 days for MGG152, BT054 and HK157; 8 days for TS603; 9 days for HK213; 11 days for HK211 and HK252).

### Metabolite quantification by GC-MS

Quantification and analysis of all metabolite levels by GC-MS (steady state levels from HOG cells, GSCs, and tissues) was performed as previously described (McBrayer et al., 2018). Briefly, to quantify metabolites in HOG cells, cells were plated in 6-well plates (0.5 × 10^6^ cells per well), washed with ice-cold saline, and snap frozen in liquid nitrogen. To extract metabolites, 350 μL 70% methanol at −20°C was added to each well, and adherent HOG cells were scraped from each well into the methanol suspension and transferred to Eppendorf tubes. Chloroform (−20°C, 150 μL) was added, and each sample was vortexed for 20 min at 4°C and centrifuged (17,000 × g for 10 min at 4°C). The upper phase containing polar metabolites in methanol was dried using a vacuum concentrator (CentriVap, Labconco) overnight at 4°C, and dried samples were stored at −80°C if not immediately used for GC-MS analysis.

To quantify metabolites from GSC lines, neurospheres were harvested from 24-well plates, followed by the addition of 4°C saline to quench metabolic activity. Samples were transferred to Eppendorf tubes, centrifuged in an Eppendorf microcentrifuge for 1 minute at 4°C, and the resulting supernatant was aspirated, with remaining cell pellets snap frozen in liquid nitrogen and stored at −80°C. Subsequent polar extraction of metabolites from cell pellets was performed as described above for HOG cells.

To quantify metabolites from tumor tissue samples, samples were homogenized using a TissueLyser II (Qiagen) with 70% methanol at −20°C (volume dependent on tissue volume). Chloroform (−20°C, volume dependent on tissue volume) was added to each sample, and subsequent centrifugation, drying, and storage was performed as described above for HOG cells.

GC-MS analysis was performed by derivatizing dried samples and analyzing using an Agilent 7890B GC/5977A MSD system. Peak integration was conducted using the Metran software tool (Yoo et al., 2008). For relative metabolite quantification in tissue culture samples, ion counts were normalized to the total ion counts detected in the sample.

### Metabolite quantification by LC-MS/MS or LC-MS

For LC-MS and LC-MS/MS analyses of tissue samples, samples were homogenized using a TissueLyser II (Qiagen) with 80% methanol at −20°C (volume dependent on tissue volume), vortexed for 20 min at 4°C, and centrifuged (21,100 × g at for 10 min at 4°C). Metabolite samples were dried using a CentriVap (Labconco) or SpeedVac (Thermo Fisher) concentrator.

For LC-MS analyses of cultured cells, cells were harvested as described above for GC-MS analysis. Metabolites were extracted in 80% methanol, vortexed for 20 min at 4°C, and centrifuged (21,100 × g for 10 min at 4°C). Metabolite samples were dried using a CentriVap (Labconco) or SpeedVac (Thermo Fisher) concentrator.

For LC-MS/MS analyses, dried metabolites were re-suspended in 20 μL HPLC-grade water. 10 μL was injected and analyzed using a 5500 QTRAP hybrid triple quadrupole mass spectrometer (AB/SCIEX) coupled to a Prominence UFLC HPLC system (Shimadzu) with selected reaction monitoring (SRM) with positive/negative polarity switching to detect a total of 263 water-soluble metabolites (Yuan et al., 2012). Amide HILIC chromatography (Waters) at pH 9.0 was used for metabolite separation over a 15 minute gradient. Peak areas were integrated using MultiQuant 2.1 software and metabolite quantification was performed using Matlab.

For LC-MS analyses, dried metabolites were resuspended in 40-50 μL 80% acetonitrile, vortexed for 20 min at 4°C, and centrifuged (21,100 × g for 10 min at 4°C). 10–20 μL was injected and analyzed with a Q-Exactive HF-X or Orbitrap Exploris hybrid quadrupole-orbitrap mass spectrometer (Thermo Fisher) coupled to a Vanquish Flex UHPLC system (Thermo Fisher). Chromatographic resolution of metabolites was achieved using on a Millipore ZIC-pHILIC column using a linear gradient of 10 mM ammonium formate pH 9.8 and acetonitrile. Spectra were acquired with a resolving power of either 120,000 or 240,000 full width at half maximum (FWHM), a scan range set to 80–1,200 m/z, and polarity switching. Data-dependent MS/MS data was acquired on unlabeled pooled samples to confirm metabolite IDs when necessary. Peaks were integrated using El-Maven 0.12.0 software (Elucidata) or TraceFinder 5.1 SP2 software (Thermo Fisher). Total ion counts were quantified using Freestyle 1.7 SP1 software (Thermo Fisher). Peaks were normalized to total ion counts using the R statistical programming language. For stable isotope tracing studies, correction for natural abundance of metabolite labeling was performed using the AccuCor package (version 0.2.3) in the R statistical programming language (Su et al., 2017).

### Absolute quantification of 2HG

Absolute 2HG quantification was performed from cell culture and tumor models using GC-MS as previously described (McBrayer et al., 2018). At the time of cell harvest for GC-MS sample preparation, cell counts and average cellular diameter values were determined in parallel cell cultures for each cell line using a Vi-CELL XR cell viability counter (Beckman Coulter). Cell number and average cellular diameter values were then used to calculate cell volume using the following formula: 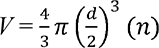, where *d* is average cellular diameter, *n* is cell number, and *V* is sample volume. For tissue samples, volumes were calculated using tissue weights and a previously published value of brain tissue density (Bothe et al., 1984). Cell and tissue samples were processed as described above in the GC-MS methods section, with 2HG concentrations calculated by dividing 2HG content by tumor sample volume. 2HG content was determined by comparison to standards containing known quantities of pure (*R*)-2HG (20 ng – 6 μg range).

### BAY 2402234 pharmacokinetics and pharmacodynamics

Naïve ICR SCID mice or mice bearing orthotopic xenografts or allografts were treated with BAY 2402234 (4 mg/kg PO QD) for 3 days. 4 hours after the last dose, mice were euthanized. Tissues were collected and processed for LC-MS or MALDI-MSI analysis.

### MALDI-MSI tissue preparation and microscopy

Intact MGG152 tumor-containing mouse brains were dissected, snap-frozen in liquid nitrogen, and stored at −80°C for MALDI-MSI analysis. The brains were cryo-sectioned in the coronal plane to 10 µm thickness and thaw-mounted onto indium tin oxide (ITO) slides. Serial sections were collected for H&E staining, and imaged using a 10× objective (Zeiss Observer Z.1, Oberkochen, Germany). A tissue drug mimetic was prepared with healthy mouse brain tissue homogenate spiked with BAY 2402234 concentrations ranging from 0.2-50 µM. The spiked homogenate was pipetted into the channels of a tissue microarray array (TMA) mold and frozen. This tissue drug mimetic was cryo-sectioned and thaw mounted adjacent to MGG152 mouse brain tissue sections.

### MALDI matrix preparation

BAY 2402234 quantitation was performed using a 2,5-dihydroxybenzoic acid (160 mg/mL) matrix solution which was dissolved in 70:30 methanol: 0.1% TFA with 1 % DMSO. The matrix was applied onto tissue using a TM sprayer (HTX Technologies, Chapel Hill, NC) with a two-pass cycle at a flow rate (0.18 mL/min), spray nozzle velocity (1200 mm/min), nitrogen gas pressure (10 psi), spray nozzle temperature (75 °C), and track spacing (2 mm). Matrix was recrystallized by incubation at 85^0^C in presence of 5% acetic acid solution. Orotate, carbamoyl aspartate, thymidine triphosphate (dTTP) and deoxycytidine triphosphate (dCTP) were imaged using a 1,5-diaminonaphthalene hydrochloride (4.3 mg/mL) matrix solution prepared in 4.5/5/0.5 HPLC grade water/ethanol/1 M HCl (v/v/v). The matrix was applied using a four-pass cycle with 0.9 mL/min flow rate, spray nozzle velocity (1200 mm/min), spray nozzle temperature (75°C), nitrogen gas pressure (10 psi), and track spacing (2 mm).

### BAY 2402234 MALDI MRM MSI

BAY 2402234 was quantitatively imaged using a timsTOF fleX mass spectrometer (Bruker Daltonics, Billerica, MA) operating in positive ion mode with a multiple reaction monitoring (MRM) method. The mass range was selected between *m/z* 100-650. The MRM settings were adjusted using a BAY 2402234-infused solution through the ESI source for the ion transfer funnels, quadrupole, collision cell, and focus pre-TOF parameters. The optimal collision energy for the BAY 2402234 precursor was 35 eV with a 3 *m/z* isolation width for the precursor to product ion transition 521.101 → 376.091 corresponding to [C_21_H_18_ClF_5_N_4_O_4_+H]^+^ and [C_15_H_12_F_4_N_3_O_4_+H]^+^ respectively. The optimized ESI method was transferred to a MALDI source method and calibrated using a tune mix solution (Agilent Technologies, Santa Clara, CA). MALDI-MSI conditions included a 10,000 Hz laser repetition rate and a 50 µm pixel size consisting of 1,000 laser shots. SCiLS Lab software (version 2021a premium, Bruker Daltonics, Billerica, MA) was used for data analysis without data normalization. The ion intensity was correlated to BAY 2402234 concentration using a linear regression between 0.2-1.0 µM from the tissue drug mimetic resulting in a correlation coefficient of 0.995. A limit of detection (LOD) of 0.10 µM (S/N ratio of > 3) and limit of quantification (LOQ) of 0.33 µM (S/N ratio of > 10) were calculated.

### Metabolite MALDI MRM MSI

Orotate, carbamoyl aspartate, dTTP, and dCTP were imaged from serial tissue sections. The Q-TOF instrument was operated in negative ion mode in full scan mode for *m/z* 50-1000. Using standards, the orotate and carbamoyl aspartate [M-H]^-^ ions fragmented in MS mode, thus the product ions were directly monitored in MS mode by setting the collision energy to 10 eV. The product ion monitored for orotate was *m/z* 111.019 and for carbamoyl aspartate *m/z* 132.029. An MRM approach was used to image dTTP (480.982→158.924) and dCTP (465.982→158.924). Further peak annotation was confirmed by performing on tissue MSMS compared to standards under the same analytical conditions.

### ^15^N tracing studies

For ^15^N stable isotope tracing studies in GSC lines, cells were cultured in human plasma-like medium (HPLM) (Cantor et al., 2017) without glutamate supplemented with 1x B27, 0.25x N2, 1% penicillin/streptomycin, amphotericin B (250 ng/mL), Plasmocin (2.5 μg/mL), EGF (20 ng/mL), bFGF (20 ng/mL), and heparin (2 μg/mL), and lacking glutamine (for amide-^15^Nglutamine glutamine tracing), uracil (for ^15^N_2_-uracil tracing), uridine (for ^15^N_2_-uridine tracing), or hypoxanthine (for ^15^N_4_-hypoxanthine tracing). Cells were acclimated to HPLM over 3 days prior to tracing (100% standard media Day 1, 50% HPLM in standard media Day 2, 100% HPLM Day 3). Tracers were added at their native HPLM concentrations: 550 µM amide-^15^N-glutamine (Cambridge Isotope Laboratories NLM-557), 3.7 µM ^15^N_2_-uracil (Cambridge Isotope Laboratories NLM-637), 3 µM ^15^N_2_-uridine (Cambridge Isotope Laboratories NLM-812), or 10 µM ^15^N_4_-hypoxanthine (Cambridge Isotope Laboratories NLM-8500).

### Metabolomics analysis of GSCs in HPLM

BT054 GSCs were cultured in HPLM without glutamate supplemented with 1x B27, 0.25x N2, 1% penicillin/streptomycin, amphotericin B (250 ng/mL), Plasmocin (2.5 μg/mL), EGF (20 ng/mL), bFGF (20 ng/mL), and heparin (2 μg/mL). Cells were acclimated to HPLM over 3 days prior to treatment (100% standard media Day 1, 50% HPLM in standard media Day 2, 100% HPLM Day 3). 24 hours after completing acclimation, media was refreshed and DMSO, 10 nM of BAY 2402234, or 5 μM lometrexol was added. Cells were harvested 24 hours after drug treatment, and metabolites were quantified by LC-MS.

### TCGA bioinformatics analysis

The TCGA Lower Grade Glioma (LGG) and Glioblastoma datasets were used to assess the genetic alterations that co-occur or are mutually exclusive with *IDH1* mutations in human gliomas (https://www.cbioportal.org). The LGG dataset was also used to generate the survival curves in patients with IDH1 WT gliomas and patients with IDH1 mutant gliomas with or without *PIK3R1*/*PIK3CA* mutations.

### Immunoblot analysis of protein expression

Cells were lysed in EBC lysis buffer containing a protease inhibitor cocktail (Sigma Aldrich 11836153001) and, for phosphoprotein immunoblots, a phosphatase inhibitor cocktail (Sigma Aldrich 04906837001). Lysates were resolved by SDS-PAGE and transferred to nitrocellulose membranes (Bio-Rad). Primary antibodies used included: anti-Akt (Cell Signaling 2920S, mouse monoclonal), anti-pAkt (Cell signaling 2965S, rabbit monoclonal), anti-GFAP (Abcam 7260, rabbit polyclonal), anti-Sox2 (Abcam 97959, rabbit polyclonal), anti-Olig2 (DF308, a gift from J. Alberta at DFCI, rabbit polyclonal) (Ligon et al., 2004), anti-nestin (Abcam 6142, mouse monoclonal), anti-γH2A.X (Cell Signaling 9718S, rabbit monoclonal), anti-HA (Covance MMS-101P, mouse monoclonal), anti-FLAG (Sigma F1804, mouse monoclonal), and anti-vinculin (Sigma V9131, mouse monoclonal). Secondary antibodies used included: Goat anti-Mouse IgG (Thermo Fisher 31430, goat polyclonal) and Goat anti-Rabbit IgG (Thermo Fisher 31460, goat polyclonal).

### Soft agar colony formation assays

Soft agar assays were performed by first plating a base layer (2.5 mL per well of a 6-well plate) of liquefied 1% agarose in DMEM medium with 10% FBS and 1% penicillin/streptomycin. Once solidified, a 1 mL mixture of 5,000 NHA cells and 0.4% agarose in DMEM medium with 10% FBS and 1% penicillin/streptomycin (warmed to 42°C) was added on top of the base layer. Once solidified, 2 mL of DMEM medium with 10% FBS and 1% penicillin/streptomycin was added to each well. Media was exchanged twice weekly. After 3 weeks, cells were stained with 200 μL of 0.1% iodonitrotetrazolium chloride (Sigma-Aldrich) and incubated overnight. Colonies were photographed and analyzed using ImageQuant TL software (GE Healthcare).

### Magnetic resonance imaging

MRI was performed using a Bruker BioSpec 7T/30 cm USR horizontal bore Superconducting Magnet System (Bruker Corp.). This system provides a maximum gradient amplitude of 440 mT/m and slew rate of 3,440 T/m/s and uses a 23 mm ID birdcage volume radiofrequency (RF) coil for both RF excitation and receiving. Mice were anesthetized with 1.5% isoflurane mixed with 2 L/min air flow and positioned on the treatment table using the Bruker AutoPac with laser positioning. Body temperature of the mice was maintained at 37°C using a warm air fan while on the treatment table, and respiration and body temperature were monitored and regulated using the SAII (Sa Instruments) monitoring and gating system, model 1025T. T2 weighted images of the brain were obtained using a fast spin echo (RARE) sequence with fat suppression. The following parameters were used for image acquisition: repetition time (TR) = 6,000 ms, echo time (TE) = 36 ms, field of view (FOV) = 19.2 × 19.2 mm^2^, matrix size = 192 × 192, spatial resolution = 100 × 100 μm^2^, slice thickness = 0.5 mm, number of slices = 36, rare factor = 16, number of averages = 8, and total acquisition time 7:30 min. Bruker Paravision 6.0.1 software was used for MRI data acquisition, and tumor volume was determined from MRI images processed using a semiautomatic segmentation analysis software (ClinicalVolumes).

### Histopathology and immunohistochemistry studies

Histopathological analysis of brain and tumor tissues was performed by harvesting tumor tissues and fixing immediately for 24 hours in 10% formalin in PBS. Following fixation, tissues where washed and stored in 70% ethanol. Tissues were embedded in paraffin, sectioned, and stained with hematoxylin and eosin (H&E). H&E sections of brain tumors were reviewed by a board-certified neuropathologist (K.L.L.) and the Dana-Farber/Harvard Cancer Center Rodent Histopathology Core.

For IHC analyses of GFAP and Olig2, 4 μm-thick tissue sections were prepared from formalin-fixed paraffin-embedded tissue blocks and air dried overnight. Slides were baked in an Isotemp Oven (Fisher Scientific) for 30–60 min at 60°C to melt excess paraffin and rehydrated. GFAP IHC was performed on the Bond III autostainer (Leica Biosystems) using the Bond Polymer Refine Detection Kit (Leica Biosystems DS9800), according to the manufacturer’s protocol. Antigens were retrieved with Bond Epitope Retrieval Solution 1 (citrate, pH 6.0) for 20 minutes. Bound antibody was visualized with a 3,3’-diaminobenzidine (DAB) stain, and counterstaining was performed with hematoxylin. Olig2 IHC was performed manually using a humidified chamber. Antigens were retrieved by heating slides with a pressure cooker to 125°C for 30 s and 90°C for 10 s in citrate buffer (pH 6; Life Technologies, 005000). Slides were then incubated with peroxidase (Dako, S2003) and protein blocking reagents (Dako, X0909), respectively, for 5 minutes each. Sections were then incubated with primary antibody, washed, and incubated with Envision+System-HRP Labeled Polymer Anti-Rabbit (Dako, K4003) for 30 minutes. Bound antibody was visualized with a 3,3’-diaminobenzidine (DAB) stain and counterstaining was performed with hematoxylin.

Primary antibodies used were anti-GFAP (Abcam 7260, rabbit polyclonal, 1:3,000) and anti-Olig2 (DF308, a gift from J. Alberta at DFCI, rabbit polyclonal, 1:20,000). The specificity of the GFAP IHC assay was determined by comparing staining in sarcoma (negative control) and normal brain (positive control) mouse tissues. The specificity of the Olig2 IHC assay was determined by comparing staining in liver (negative control) and normal brain (positive control) mouse tissues.

IHC analyses of γH2A.X and cleaved caspase 3 were performed by Histowiz, Inc. (histowiz.com) using a Standard Operating Procedure and fully automated workflow. Samples were processed, embedded in paraffin, and sectioned at 4 μm. Immunohistochemistry was performed on a Bond Rx autostainer (Leica Biosystems) with Heat Induced Epitope Retrieval (HIER). Antibodies used were: anti-γH2A.X (Cell Signaling 9718, rabbit polyclonal, 1:800) and anti-cleaved caspase 3 (Cell Signaling 9661, rabbit monoclonal, 1:300). Chromagen development was done using the Bond Polymer Refine Detection Kit (Leica Biosystems) which was used according to the manufacturer’s protocol. After staining, sections were dehydrated and film coverslipped using a TissueTek-Prisma Coverslipper. Whole slide scanning (40x) was performed on an Aperio AT2 (Leica Biosystems). To quantify cleaved caspase 3 and γH2A.X staining, regions of interest (ROIs) were defined and total and chromogen-positive cells were counted using QuPath software (version 0.2.3). For CC3 quantification, 12–16 ROIs were analyzed from 2 sections per organoid. For γH2A.X quantification, 16 ROIs were analyzed from 2 vehicle-treated tumors and 86 ROIs were analyzed from 5 BAY 2402234-treated tumors. Spectral dissociation was used to separate signals from either the chromogen (DAB) and the counterstain (hematoxylin).

#### Validation of genetic alterations in engineered astrocytomas

For validation of genetic alterations in our GEM model and in DF-AA27 GSCs, RNA and genomic DNA were harvested from astrocytoma tissue, DF-AA27 cells, normal mouse brain tissue, and genetically engineered MEFs. Genomic DNA was extracted using the DNeasy Blood and Tissue kit (Qiagen) per manufacturer’s instructions. To assess CRISPR-based editing of *Atrx* and *Trp53*, nested PCR was performed using genomic DNA templates to generate amplicons centered on the PAM sites targeted by *Atrx* and *Trp53* sgRNAs. The KOD Xtreme polymerase kit (EMD Millipore 71975) was used for PCR. For *Trp53*, the following set of primers were used in sequential PCR reactions to generate amplicons for sequencing:

Outer forward *Trp53:* ATAGAGACGCTGAGTCCGGTTC
Outer reverse *Trp53*: CCTAAGCCCAAGAGGAAACAGA
Inner forward *Trp53:* TGCAGGTCACCTGTAGTGAGGTAGG
Inner reverse *Trp53:* GAAACAGGCAGAAGCTGGGGAAGAAAC

The inner PCR reaction was repeated using the above inner primers for the third and final PCR reaction for *Trp53*.

For *Atrx*, the following set of primers were used in sequential PCR reactions to generate amplicons for sequencing:

Outer forward *ATRX:* GCTATCTGAAACTCAATCCACG
Outer reverse *ATRX:* GACTTGGTTTCTCCTTTGCCATG
Inner forward *ATRX:* GCTTCCTGTAAGCTCATAAGTAC
Inner reverse *ATRX*: CTAATGCCATATGAGTGTAACTC
Second inner forward *ATRX*: CTCTTACATAATGGCCATTCTC
Second inner reverse *ATRX*: CTGTGAGTCATGATCATTCTTTGC

PCR products were purified using the NEB Monarch kit (T1030S) between each PCR reaction. Final PCR products were gel-purified using the Qiagen Gel Extraction Kit (Qiagen 28706) and submitted for next-generation sequencing using the MGH CCIB DNA Core.

Presence of the *Idh1-R132H* mutation was assessed using a PCR assay specific for the recombined lox-STOP-lox (LSL) element as previously described (Sasaki et al., 2012b). Nested PCR amplification was performed using genomic DNA templates with primers for either the recombined *Idh1-R132H* allele or *Actb* (encoding beta actin):

Forward primer recombined *Idh1-R132H:* GATTGATTCTGCCGCCATGATCCTAGT
Reverse primer recombined *Idh1-R132H:* CCTGGTCATTGGTGGCATCACGATTCTC
Forward primer *Actb*: TGACCCAGGTCAGTATCCCGGGT
Reverse primer *Actb*: GAACACAGCTAAGTTCAGTGTGCTGGGA

Expression of the *Pik3ca-H1047R* allele was assessed by harvesting RNA and performing RT-PCR as previously described (Adams et al., 2011). RNA was isolated using the RNeasy Mini Kit (Qiagen 74106) prior to cDNA synthesis using the AffinityScript qPCR cDNA Synthesis Kit (Agilent 600559). PCR was performed, and the following primers were used to detect the expression of the *Pik3ca-H1047R* transgene or *Actb*:

Forward primer *Pik3ca-H1047R*: CTAGGTAGGGGATCGGGACTCT
Reverse primer *Pik3ca-H1047R*: AATTTCTCGATTGAGGATCTTTTCT
Forward primer *Actb*: TAGGCACCAGGGTGTGATG
Reverse primer *Actb*: CATGGCTGGGGTGTTGAAGG

### 53BP1 foci formation assays

HOG-EV or HOG-R132H cells were plated at a density of 3 x 10^4^ cells per well onto 18 mm glass coverslips in 12-well plates. The following day, cells were treated with either DMSO or 10 nM BAY 2402234. Cells were fixed at the indicated time points with 4% formaldehyde, permeabilized with 0.3% Triton X-100 in PBS, incubated with Triton Block (0.2 M glycine, 2.5% FBS, 0.1% Triton X-100 in PBS), and immunostained with an anti-53BP1 antibody (1:1,000, Novus Biologicals, NB100-304) diluted in Triton Block. Cells were washed, incubated with a donkey anti-rabbit Alexa Fluor 488 secondary antibody (Invitrogen, A21206), and mounted. Images were acquired using a DeltaVision Ultra (Cytiva) microscope equipped with a 60x objective with 9 x 0.5 μm z-sections. Images were deconvolved and maximum intensity projections were generated. 53BP1 foci were quantified in individual cells using ImageJ with a macros plugin (*n* = 325–505 cells per condition per time point).

### FACE quantification of lipid-linked oligosaccharides

FACE assays were conducted as previously described (Gao et al., 2013). Briefly, HOG-EV or HOG-R132H cells were plated at a density of 2.5 x 10^6^ cells in 15 cm dishes. 24 hours later, 10 nM BAY 2402234, 1 μg/mL tunicamycin, or DMSO was added. 24 hours later, plates were transferred to wet ice, medium was aspirated, and cells were rinsed twice with 4°C PBS. 12 mL methanol at room temperature was added to each plate and cells were lifted from plates using cell scrapers and transferred to 15 mL conical tubes. Lipid-linked oligosaccharides (LLOs) were recovered from methanolic cell suspensions and subjected to hydrolysis in weak acid. Oligosaccharides were labeled with the fluorophore 7-amino-1,3-naphthalenedisulfonic acid (ANDS) and resolved by electrophoresis. Fluorophore-labeled LLOs and oligosaccharide standards were detected with a fluorescence imager.

### dC and dT supplementation experiments

HOG-EV or HOG-R132H cells were plated at a density of 1 x 10^5^ cells per well in 6-well plates (for cell death analysis), 2.5 x 10^5^ cells per well in 6-well plates (for metabolomics), or at 6 x 10^5^ cells in 10 cm dishes (for γH2A.X immunoblots). 24 hours later, BAY 2402234 was added with or without 15 μM dC, 15 μM dT, or both. Cells were harvested after 72 hours (cell death analysis), 24 hours (for metabolomics), or 48 hours (for γH2A.X immunoblots).

### Palbociclib rescue experiments

HOG-EV or HOG-R132H cells were pre-treated for 24 hours with 1 μM Palbociclib or DMSO in 10 cm dishes. Next, cells were harvested, counted and plated at a density of 1 x 10^5^ cells per well in 6-well plates (for cell death analysis) or at 6 x 105 cells in 10 cm dishes (for γH2A.X immunoblots) and 1 μM Palbociclib or DMSO was re-added. 24 hours later, BAY 2402234 was added. Cells were harvested 72 hours or 36 hours later for cell death analysis or γH2A.X immunoblots, respectively.

### Acute (*R*)-2HG manipulation experiments

Acute manipulation of intracellular (*R*)-2HG levels during DHODH inhibition involved pre-treatment/concurrent treatment with AGI-5198 or (*R*)-2HG-TFMB. DF-AA27 GSCs were treated with 3 μM AGI-5198 for 72 hours prior to being harvested for 2HG quantification by GC-MS. HOG-R132H cells were pretreated for 72 hours with 3 μM AGI-5198. Then media was exchanged with media containing 3 μM AGI-5198 and 2 mL of IMDM with 500 nM or 1,000 nM brequinar. 48 hours later, cells were washed with PBS, trypsinized, and harvested for cell death quantification. 50,000 MGG152 cells were plated per well in a 24-well low-adherent plate in 250 μL media with 110 nM AGI-5198. After 48 hours of AGI-5198 pre-treatment, 250 μL media was added to each well with 2x the indicated concentration of brequinar and 110 nM AGI-5198. 96 hours later, cells were washed with PBS, trypsinized, and harvested for cell death quantification. HOG-EV cells were plated at a density of 1.5 × 10^5^ cells per well in 6-well plates with 2 mL of IMDM media per well. After 24 hours, the media was exchanged with media containing stocks of DMSO or (*R*)-2HG-TFMB at the indicated concentrations. After 3 hours, 5 nM BAY 2402234 was added to each well. After 24 hours, media was replenished with (*R*)-2HG-TFMB and BAY 2402234 at the same doses. 24 hours later, cells were washed with PBS, trypsinized, and harvested for cell death quantification.

## QUANTIFICATION AND STATISTICAL ANALYSIS

Information related to data presentation and statistical analysis for individual experiments can be found in the corresponding figure legends. Statistical analyses were carried out using GraphPad Prism software or the web-based metabolomics data analysis tool MetaboAnalyst. Significance of all comparisons involving two groups was calculated by unpaired two-tailed *t*-test. For comparisons of two groups with significantly different variances, Welch’s *t*-test was used. For comparisons of two groups without significant differences in variances, Student’s *t*-test was used. Significance of all comparisons involving three or more groups was calculated by one-way ANOVA. Significance of survival data was determined by log-rank test. Statistical evaluation of patterns of co-occurrence or mutual exclusivity were performed using one-sided Fisher’s exact tests. Significance of correlations between 2HG content and drug sensitivity was determined by simple linear regression analysis. For all tests, *p-*values < 0.05 were considered statistically significant.

