## Supplemental Table 2 for "De Novo Pyrimidine Synthesis is a Targetable Vulnerability in IDH Mutant Glioma"

| **Table S2. Genetic Alterations Co-Occurring and Mutually Exclusive with IDH1 Mutations in Glioma** | | | | | | |
| --- | --- | --- | --- | --- | --- | --- |
|  |  |  | Lower Grade Glioma  (TCGA, Provisional) | | Glioblastoma  (TCGA, Provisional) | |
| Gene 1 | Gene 2 | Association | p-Value | Log Odds Ratio | p-Value | Log Odds Ratio |
| IDH1 | TP53 | Co-Occurrence | < 0.001 | > 3.00 | < 0.001 | > 3.00 |
| IDH1 | ATRX | Co-Occurrence | < 0.001 | > 3.00 | < 0.001 | > 3.00 |
| IDH1 | CIC | Co-Occurrence | < 0.001 | 2.155 | 0.048 | > 3.00 |
| IDH1 | PIK3R1 | Co-Occurrence* | 0.134 | 1.991 | 0.011 | 2.431 |
| IDH1 | PIK3CA | None | 0.411 | -0.289 | 0.536 | -0.700 |
| IDH1 | PTEN | Mutual Exclusivity | < 0.001 | < -3.00 | < 0.001 | < -3.00 |

*Statistically significant co-occurrence observed only in glioblastoma study.
