## Supplementary figures and images for "De Novo Pyrimidine Synthesis is a Targetable Vulnerability in IDH Mutant Glioma"

### Figure S1

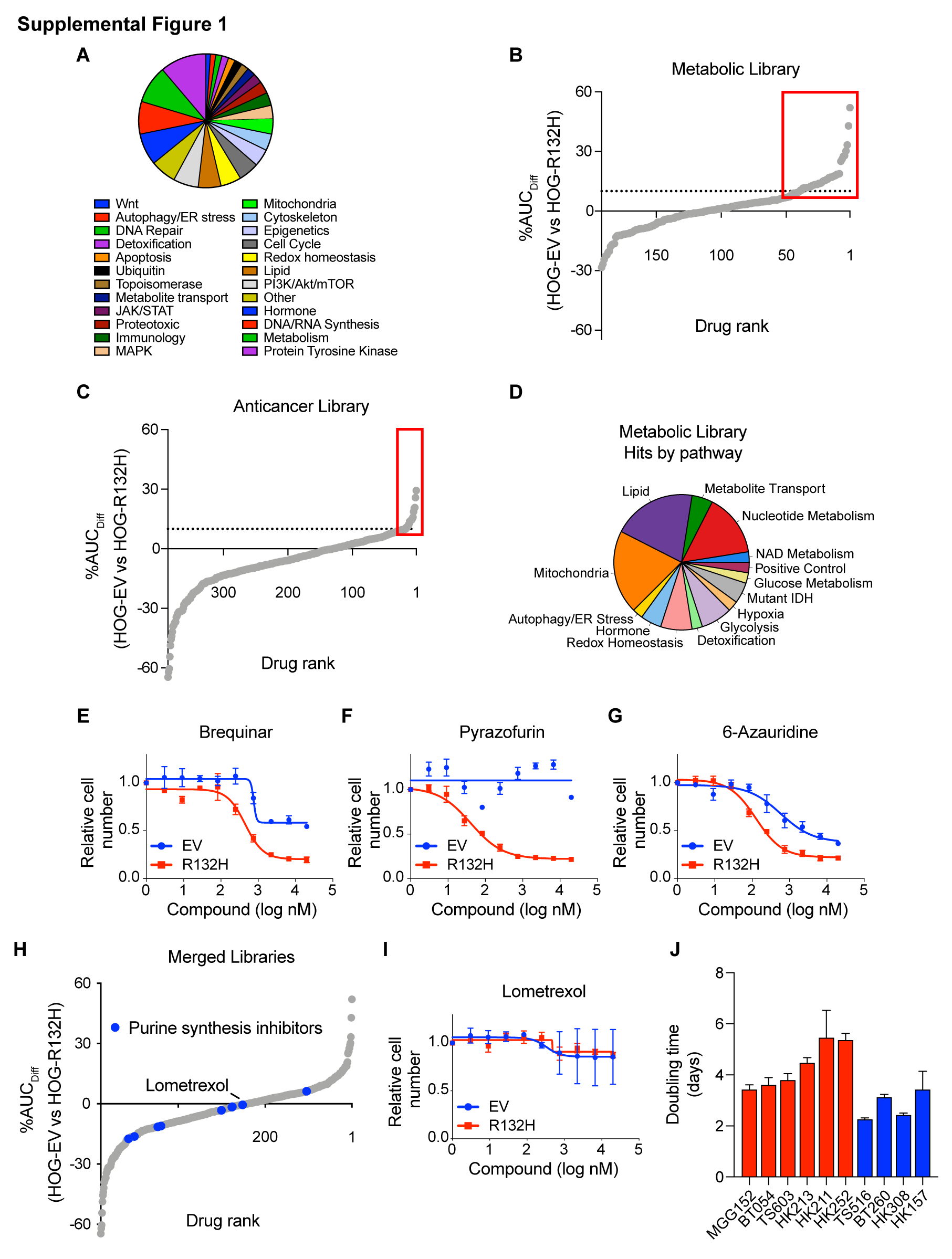

### Figure S2

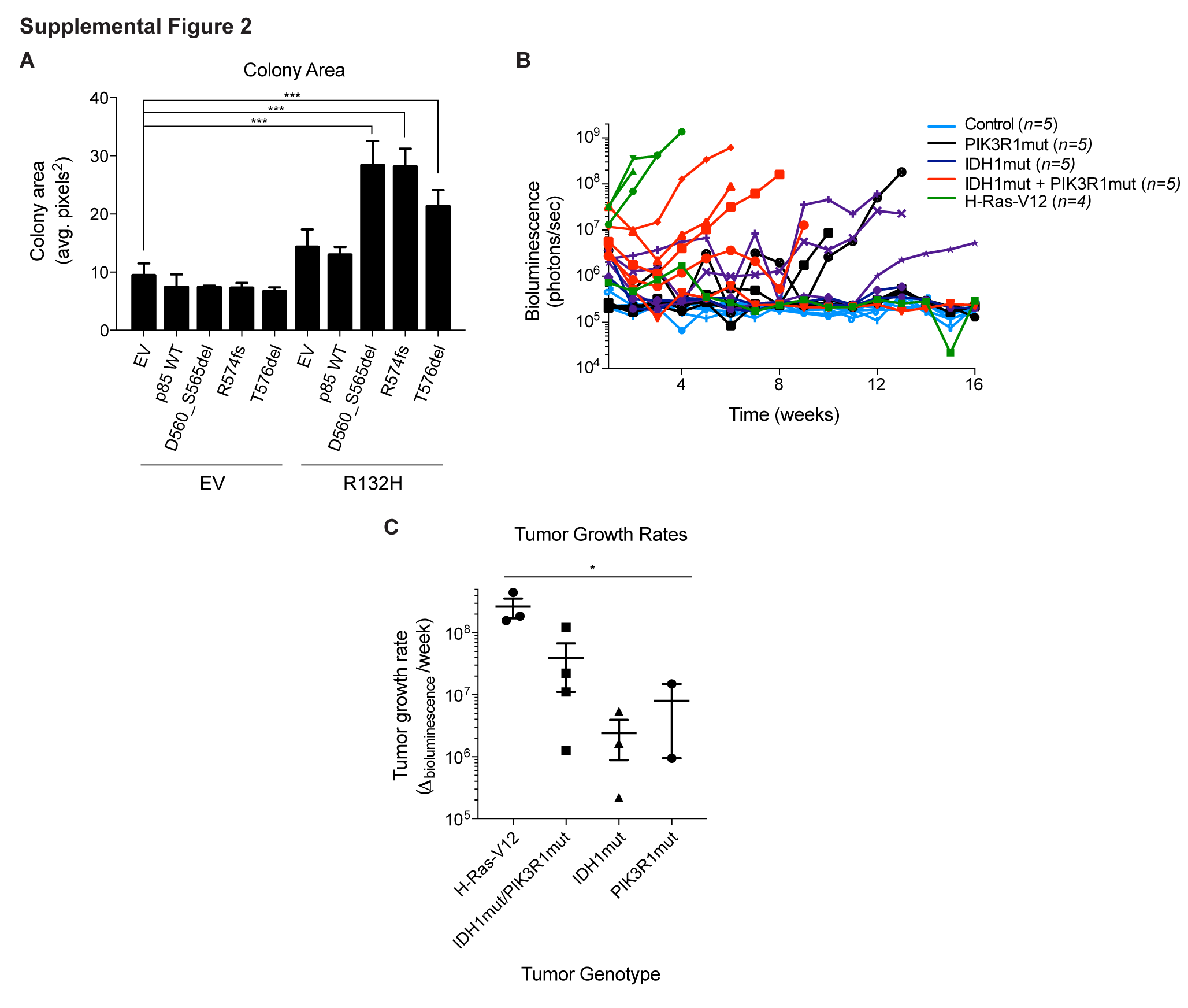

### Figure S3

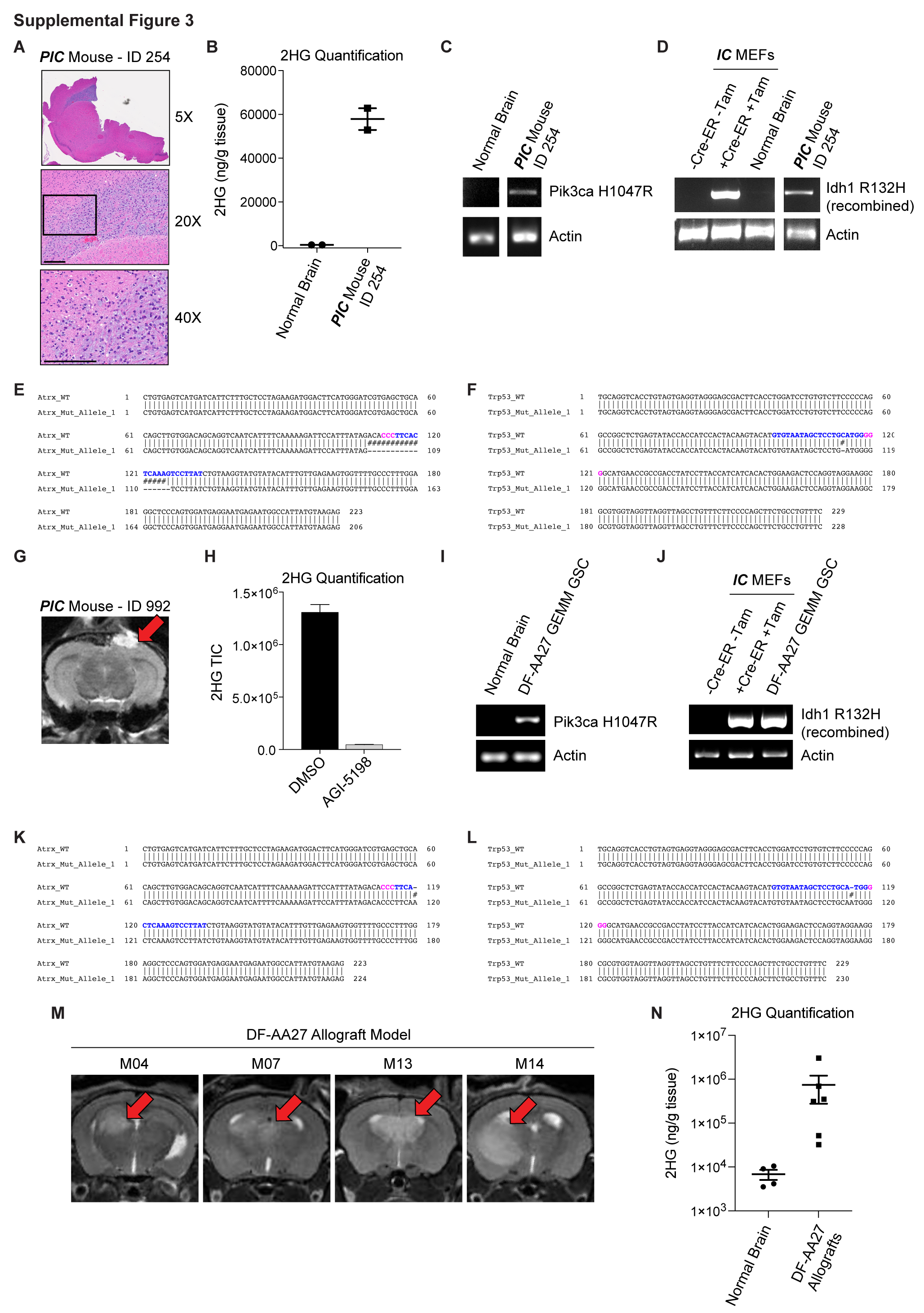

### Figure S4

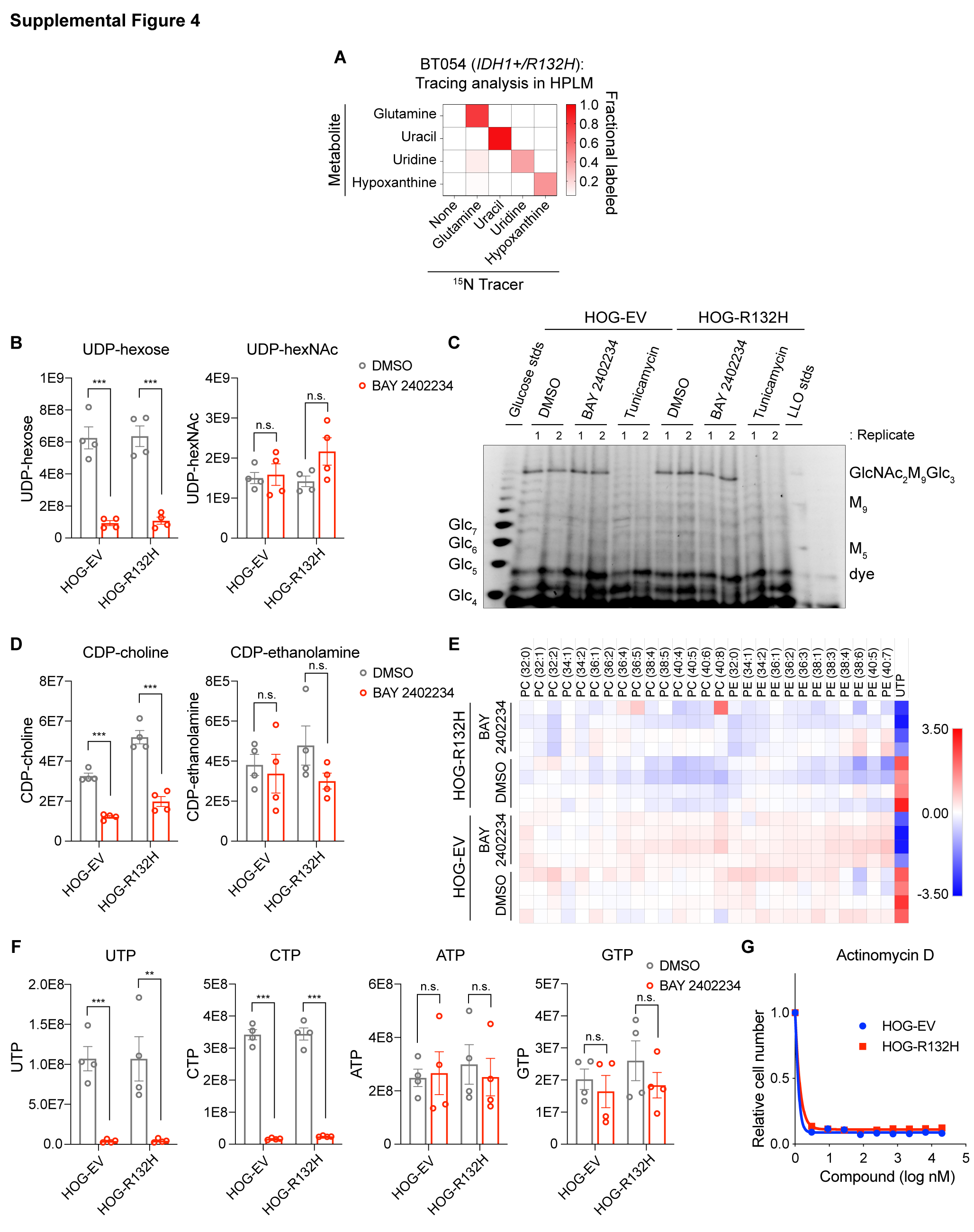

### Figure S5

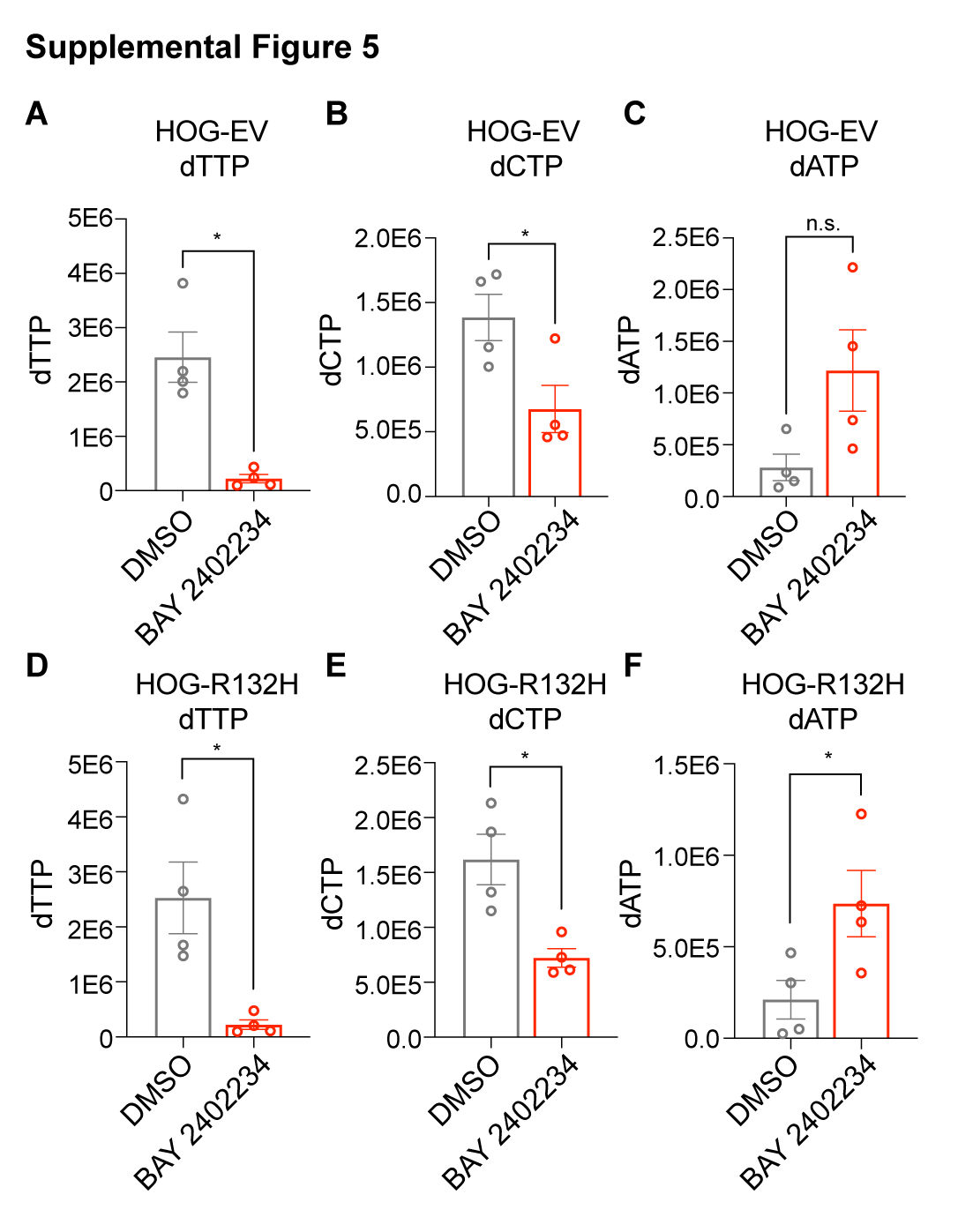

### Figure S6

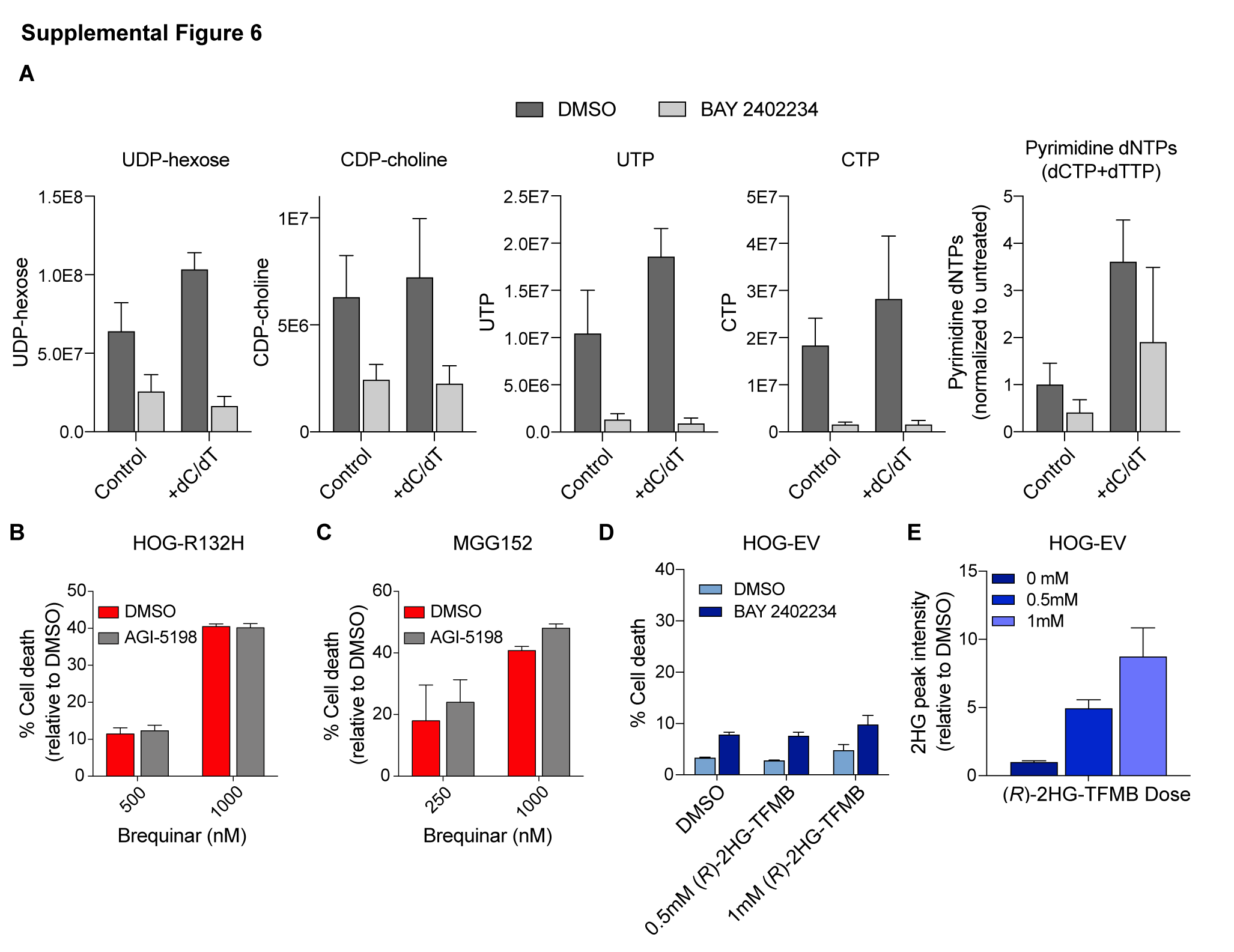
