## Supplemental Table Legends for "De Novo Pyrimidine Synthesis is a Targetable Vulnerability in IDH Mutant Glioma"

**Table S1. Results of Combined Metabolic Inhibitor Library and Anticancer Drug Library Screen, Related to Figure 1.** For each drug, relative cell number versus drug dose was plotted for HOG-EV and HOG-R132H cells and area under the curve (AUC) values were calculated. %AUC_Diff_ values in HOG-EV and HOG-R132H cells were calculated for all drugs, which were then used to rank drugs according to their preferential activity against IDH1 mutant glioma cells.

**Table S2. Genetic Alterations Co-Occurring and Mutually Exclusive with IDH1 Mutations in Glioma, Related to Figure 4.** Analysis of associations between genetic alterations in *IDH1* and *TP53, ATRX, CIC, PIK3R1, PIK3CA,* and *PTEN* from The Cancer Genome Atlas (TCGA) Lower Grade Glioma (*n =* 283) and Glioblastoma (*n =* 585) datasets. One-sided *p*-values were determined by one-sided Fisher exact test. * indicates statistically significant co-occurrence observed only in Glioblastoma study.
