## Supplemental Figure Legends for "De Novo Pyrimidine Synthesis is a Targetable Vulnerability in IDH Mutant Glioma"

**SUPPLEMENTAL FIGURES**

**Figure S1. Drug Screen in IDH1 Mutant and IDH1 WT Glioma Cells, Related to Figure 1.** (A) Pie chart depicting pathways targeted by drugs in the metabolic and anticancer libraries. (B and C) Waterfall plot of drugs screened from the metabolic library (B) and the anticancer library (C). Hits were determined by plotting relative cell number by dose of each drug tested for both HOG-EV and HOG-R132H cells and calculating the areas under each curve (AUCs). %AUC_Diff_ values in HOG-EV and HOG-R132H cells were calculated for all drugs. Drugs with >10%AUC_Diff_ (dotted line) were categorized as hits (red outline). (D) Pie chart depicting pathways targeted by hits in the metabolic library used in (B). (E–G) Drug screen results for HOG-EV and HOG-R132H cells treated with brequinar (E), pyrazofurin (F), or 6-azauridine (G). (H) Waterfall plot of drugs from the metabolic and anticancer libraries with purine synthesis inhibitors highlighted in blue. (I) Drug screen results, as in (E–G), for lometrexol, an inhibitor of de novo purine synthesis. (J) Doubling times of IDH1 mutant (MGG152, BT054, TS603, HK213, shown in red) and IDH wild-type (TS516, BT260, HK308, HK157, shown in blue) GSC lines. For HK211, *n =* 2; for all others, *n =* 3. For all panels, data presented are means ± SEM.

**Figure S2. In Vitro and In Vivo Assessments of Transformation of Engineered Astrocytes, Related to Figure 4.** (A) Average sizes of soft agar colonies formed by the indicated NHA cell derivatives 21 days after plating. NHA cells were infected to express HA-tagged IDH1-R132H [or with the empty vector (EV)] and then superinfected to express FLAG-tagged wild-type or mutant (D560_S565del, R574fs, or T576del) p85 (protein product of *PIK3R1* gene) or the corresponding EV. All cells were also infected to express firefly luciferase and GFP. *n =* 5. (B) Quantification of brain bioluminescence after intracranial injection of astrocytes expressing EV/EV (control), IDH1^R132H^/EV (IDH1mut), EV/p85^D560_S565del^ (PIK3R1mut), IDH1^R132H^/p85^D560_S565del^ (IDH1mut + PIK3R1mut), or H-Ras-V12 (as a positive control) as well as firefly luciferase and GFP. Bioluminescence imaging was performed once per week for 16 weeks; time 0 = one week post-tumor cell injection. (C) Quantification of xenograft tumor growth rates in mice described in (B). Mice that did not develop tumors or were not able to be imaged ≥ 2 times were censored because growth rates could not be calculated in these instances. For all panels, data presented are means ± SEM; **p* < .05, **** p* < .001. In (A), two-tailed *p-*values were determined by unpaired *t*-test. In (C), *p-*value was determined by one-way ANOVA.

**Figure S3. Characterization of a GEM Model of Astrocytoma Driven by Mutant IDH1, Related to Figure 5.** (A) Representative images from hematoxylin and eosin (H&E) stained sections of brain tissue from an AAV-injected ***PIC*** mouse. Higher magnification images show area of infiltrating tumor cells into normal brain parenchyma. Black box in middle panel indicates magnified region in lower panel; scale bars = 200 μm. (B) Absolute 2HG levels (measured by GC-MS) in tumor tissue extracted from the mouse brain shown in (A) and in normal mouse brain tissue; *n* = 2. (C) RT-PCR assays for *PIK3CA^H1047R^* transgene or *Actb* gene expression in tumor tissue extracted from the mouse brain shown in (A) and in normal mouse brain tissue. (D) Genomic DNA PCR assays to detect the recombined *LSL-Idh1-R132H* allele or *Actb* gene in tumor tissue extracted from the mouse brain shown in (A), normal mouse brain tissue, and in mouse embryonic fibroblasts (MEFs) lines derived from a naïve ***IC*** mouse. MEFs were transduced in vitro with retrovirus to express the tamoxifen-inducible MerCreMer recombinase and then treated with 4-hydroxytamoxifen or were mock transduced and untreated to serve as positive and negative controls, respectively. (E and F) CRISPR amplicon sequencing of genomic DNA regions of *Atrx* (E) and *Trp53* (F) genes targeted by sgRNAs shown in Figure 5A in tumor tissue extracted from the mouse brain shown in (A). Dominant variant alleles identified in tumor tissue are aligned with wild-type gene sequences. PAM sites and sgRNA sequences are shown in pink and blue, respectively; # indicate mismatches and – indicate missing DNA bases. (G) Representative MRI image of an AAV-injected ***PIC*** mouse that developed a needle-track osteosarcoma at the injection site in the skull. (H) Relative 2HG levels (measured by GC-MS) in DF-AA27 GSCs treated with 3 μM AGI-5198 (a mutant IDH1 inhibitor) or DMSO for 72 hours; *n* = 2. TIC = total ion count. (I) RT-PCR assays for *PIK3CA^H1047R^* transgene or *Actb* gene expression, as in (C), in DF-AA27 GSCs. (J) Genomic DNA PCR assays to detect the recombined *LSL-Idh1-R132H* allele or *Actb* gene, as in (D), in DF-AA27 GSCs. (K and L) CRISPR amplicon sequencing of genomic DNA regions of *Atrx* (K) and *Trp53* (L) genes in DF-AA27 GSCs, as in (E) and (F). (M) Representative MRI images of mouse brains with DF-AA27 allografts. Unique mouse IDs are shown above each image. (N) Absolute 2HG levels (measured by GC-MS) in DF-AA27 allografts (*n =* 6) and normal mouse brain (*n =* 4). For (B) and (H), data presented are means ± standard deviation. For (N), data presented are means ± SEM.

**Figure S4. Quantification of Protein Glycosylation, Phospholipid Synthesis, and RNA Synthesis Intermediates, Related to Figure 7.** (A) ^15^N stable isotope tracing assays using the indicated ^15^N-labeled metabolites in human plasma-like medium (HPLM) (*n =* 3 per tracer). Heatmap depicts labeling of intracellular metabolite pools (y-axis) by tracers (x-axis) at 18 hours. (B), (D), and (F) Relative quantification of protein glycosylation (B), phospholipid synthesis (D), and RNA synthesis (F) intermediates by LC-MS in HOG-EV and HOG-R132H cells treated with 10 nM BAY 2402234 or DMSO for 24 hours (*n =* 4). Values on y-axes are peak areas determined by LC-MS. (C) Fluorophore-Assisted Carbohydrate Electrophoresis (FACE) assay of lipid-linked oligosaccharides (LLOs) in HOG-EV and HOG-R132H cells treated with 10 nM BAY 2402234, 1 μg/mL tunicamycin, or DMSO for 24 hours (*n =* 2). Glucose oligomer and LLO standards (stds) are shown in the first lane and the second to last lane; Glc = glucose, M = mannose, GlcNAc = N-acetylglucosamine. Dolichol-linked GlcNAc_2_M_9_Glc_3_ oligosaccharide (labeled at right) is a substrate for N-linked glycosylation and is not regulated by BAY 2402234 treatment. Tunicamycin, an inhibitor of N-linked glycosylation, depletes dolichol-linked GlcNAc_2_M_9_Glc_3_ and serves as a positive control. (E) Heat map depicting fold change of phosphatidylcholine (PC) and phosphatidylethanolamine (PE) lipids in HOG-EV and HOG-R132H cells treated with 10 nM BAY 2402234 or DMSO for 24 hours (*n =* 4). UTP is a pharmacodynamic marker for DHODH inhibition and is included as a positive control. (G) Drug screen results for HOG-EV and HOG-R132H cells treated with the mRNA synthesis inhibitor actinomycin D. Data are derived from drug screen depicted in Figure 1A–C. For all panels, data presented are means ± SEM. ***p* < .01, ****p* < .001, n.s. = not significant. Two-tailed *p*-values were determined by unpaired *t*-test.

**Figure S5. Metabolomics Analysis of Deoxynucleotide Triphosphate Levels, Related to Figure 7.** (A–F) Relative quantification of pyrimidine (dTTP, dCTP) and purine (dATP) deoxynucleotide triphosphate (dNTPs) levels by LC-MS in HOG-EV (A–C) and HOG-R132H (D–F) cells treated with 10 nM BAY 2402234 or DMSO for 24 hours (*n =* 4). Values on y-axes are peak areas determined by LC-MS. For all panels, data presented are means ± SEM; **p* < .05, n.s. = not significant. Two-tailed *p*-values were determined by unpaired *t*-test.

**Figure S6. Mechanistic Studies of De Novo Pyrimidine Synthesis Hyperdependence Induced by IDH Mutations, Related to Figure 8.** (A) Relative quantification of protein glycosylation (UDP-hexose), phospholipid synthesis (CDP-choline), and RNA synthesis (UTP, CTP) intermediates and pyrimidine dNTPs by LC-MS in HOG-R132H cells treated with 10 nM BAY 2402234 or DMSO in the presence or absence of 15 μM deoxycytidine (dC) and 15 μM deoxythymidine (dT) for 24 hours (*n =* 4). (B and C) Cell death assays of HOG-R132H (B) and MGG152 (C) cells treated with the indicated doses of brequinar for 48 and 96 hours, respectively, with or without pre-treatment with the mutant IDH inhibitor AGI-5198. HOG-R132H cells were pre-treated with 3 μM AGI-5198 for 72 hours (*n =* 3); MGG152 cells were pre-treated with 110 nM AGI-5198 for 48 hours (*n =* 2). (D) Cell death assays of HOG-EV cells treated with 5 nM BAY 2402234 or DMSO for 48 hours. Cells were pre-treated with the indicated doses of cell-permeable 2HG ester, (*R*)-2HG-TFMB, or DMSO for 3 hours and media was refreshed every 24 hours with freshly added drug, ester, and/or vehicle (*n =* 2). (E) Relative quantification of 2HG levels by GC-MS following treatment with the indicated doses of (*R*)-2HG-TFMB or DMSO for 5 hours (*n =* 2). For all panels, data presented are means ± SEM.
